# Inverse Material Parameter Estimation of Patient Specific Finite Element Models at the Carotid Bifurcation: The Impact of Excluding the Zero Pressure Configuration and Residual Stress

**DOI:** 10.1101/2022.04.12.487823

**Authors:** R. D Johnston, M. Ghasemi, C. Lally

## Abstract

The carotid bifurcation experiences a complex loading environment due to its anatomical structure. Previous *in-vivo* material parameter estimation methods often use simplified model geometries, isotropic hyperelastic constitutive equations or neglect key aspects of the vessel, such as the zero-pressure configuration or residual stress. These factors have independently been shown to alter the stress environment of the vessel wall. Characterising the location of high stress in the vessel wall has often been proposed as a potential indicator of structural weakness. However, excluding the afore-mentioned zero-pressure configuration, residual stress and patient specific material parameters can lead to an incorrect estimation of the true stress values observed, meaning stress alone as a risk indicator of rupture is insufficient. In this study, we investigate how the estimated material parameters and overall stress distributions in geometries of carotid bifurcations, extracted from *in-vivo* MR images, alter with the inclusion of the zero-pressure configuration and residual stress.

This approach consists of the following steps: (1) geometry segmentation and hexahedral meshing from *in-vivo* MRI images at two known phases; (2) computation of the zero-pressure configuration and the associated residual stresses; (3) minimisation of an objective function built on the difference between the stress states of an “ almost true” stress field at two known phases and a “deformed” stress field by altering the input material parameters to determine patient specific material properties; and (4) comparison of the stress distributions throughout these carotid bifurcations for all cases with estimated material parameters. This numerical approach provides insights into the need for estimation of both the zero-pressure configuration and residual stress for accurate material property estimation and stress analysis for the carotid bifurcation, establishing the reliability of stress as a rupture risk metric.

**Graphical Abstract:** 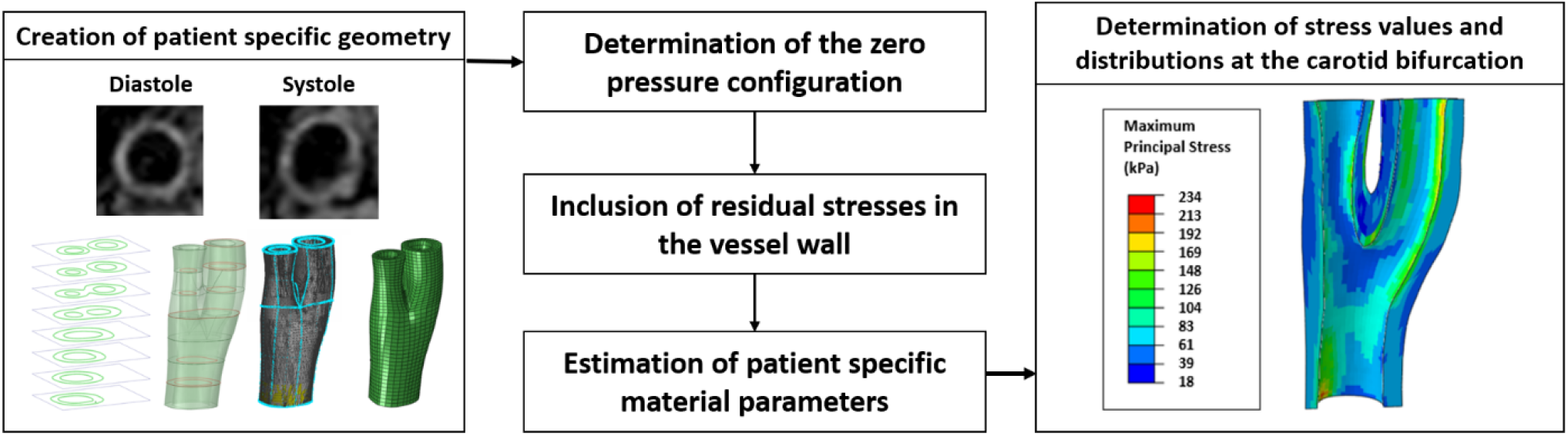

## 1. Introduction

Cardiovascular disease (CVD) is known to be the leading cause of death worldwide [1]. Currently, it is estimated that CVDs were responsible for an estimated 17.8 million deaths in 2017, more than 30% globally and this is predicted to increase in the future [1]–[3]. The most common CVDs to present themselves are aneurysms and atherosclerosis [1], both of which occur due to microstructural changes in the vessel wall. Standard diagnosis techniques are based off geometrical measures such as maximum dilation in the case of aneurysm and percent stenosis in the case of vulnerable plaques [4], [5]. Percent stenosis, a measure based on luminal narrowing, is an inadequate measure when used alone, as it does not account for the strength of the vessel wall and plaque microstructure. Unfortunately, this results in unnecessary invasive surgeries being performed due to an inability to determine vessel and plaque mechanical stability [6]. New diagnostic metrics that can aid physician treatment decisions need to be developed. Many studies have used patient specific finite element (FE) models to calculate stresses and strains for possible rupture predictions in both aneurysms and atherosclerotic plaques [7]–[16]. These models are often derived from diagnostic images such as CT [17]–[21], Ultrasound [22], [23] or MRI [24], [25] and provide physiological data that can act as inputs for computational analysis. Accurately characterising stresses and strains in patient specific models could lead to a more accurate diagnosis of vessel or plaque vulnerability when compared to the degree of vessel stenosis.

For accurate patient specific finite element (FE) analysis, robust geometry segmentation and meshing is important [20], [21], however, using commercially available finite element solvers to hexahedral mesh complex 3D arterial geometries is very difficult. To-date, some studies have created structured hexahedral meshes of carotid bifurcations [17], [18], [20], [21], [24], [26]–[29], but in many cases simplifying assumptions are used. These assumptions range from assigning a uniform wall thickness by extruding the lumen, to assuming the geometry is cylindrical or using a one-layer vessel model. Using current in-vivo imaging techniques, geometries extracted from medical images are already in a loaded configuration due to a physiological pressure being present in the artery [30]. Directly simulating this geometry will give incorrect deformations to those *in-vivo*. The zero-pressure configuration must therefore be estimated to ensure accurate simulations. Multiple studies have looked at estimating the zero pressure configuration from *in-vivo* images [30]–[34]. These methods generally involve a forward projection of the loaded *in-vivo* geometry with the corresponding pressure to determine a deformed configuration. Using a displacement calculation, a back projection of the initial geometry is then determined to estimate the zero-pressure configuration. A minimisation algorithm, usually based on displacement difference between the pressurised zero pressure configuration and the original in-vivo geometry, is then implemented to increase the accuracy of the estimated zero pressure configuration [30]–[34].

It has been well documented in the literature that residual stresses are present in arterial vessels [35]– [38]. These residual stresses have a significant impact on the distribution of physiological stresses throughout the vessel wall and tend to homogenise the stress distribution within each arterial layer in the physiological state. Numerous studies have demonstrated that the inclusion of residual stresses under physiological loading conditions substantially reduces the variation in both the circumferential and axial residual stress within the vessel wall [39]–[43]. Therefore, for structural analysis of the vessel wall, exclusion of the residual stresses in any computational model would typically lead to an over estimation of the actual stresses experienced *in-vivo*. Excluding the residual stress in the vessel will therefore reduces the validity of vessel stress being used as a vulnerability metric. To include the residual stress in FE models, the classical “opening angle” experiment is often performed [39], [40], [43]–[45]. By cutting the artery in the radial direction, the vessel springs open due to the release of the circumferential residual stresses in the vessel, which are tensile on the outer wall of the vessel and compressive on the inner wall. This opened vessel now serves as the initial configuration and can be closed to incorporate these residual stresses back into the vessel. Although these methods of implementing residual stresses are often deemed the gold standard, they are not without limitations. Depending on the location of the cut or the geometry of the vessel, the opening angle can be different, even in the same artery [39]. Furthermore, depending on age and gender, opening angles for every person would be different [46] and this cannot be determined directly *in-vivo*. Therefore, for inclusion of residual stresses in patient specific analyses, a more theoretical approach is necessary. Many studies have investigated the inclusion of residual stress numerically and without the use of the opening angle and have demonstrated their ability to replicate deformations observed *in-vivo* [47]– [49].

For material parameter estimation of arterial vessels, many studies have taken excised tissue from animal models or human cases [50]–[55]. Mechanical tests are then performed, which are either uniaxial or biaxial in nature, to determine the stress-strain response of the tissue [56]–[58]. This procedure is then simulated in FEA with estimated material parameters and then optimised until convergence is observed between experimental and simulated data [56], [59]. Although these methods provide insight into alternative material parameter estimation techniques and the specific nature of each, they are unable to be translated for material parameter estimation *in-vivo*. Therefore, alternative methods have used a combination of both *in-vivo* imaging and inverse FEA to estimate material parameters based on stresses or strains obtained [60]–[62]. Liu et al, assumed the vessel to be statically determinate such that the stress in the deformed configuration is insensitive to material properties, giving an “almost true” or “apparent” stress distribution of the vessel wall when attributed very stiff material properties [60], [63]. This method was also numerically validated and provides a robust method to estimate material parameters *in-vivo*.

In this study, we propose a new robust method for the creation and hexahedral meshing of arterial geometries from *in-vivo* MR images using available commercial software. This method can provide structured hexahedral meshes of complex arterial bifurcations and can be exported to the desired finite element solver. The method can also be extended to include the multiple layers of the vessel, plaque components and be tailored for other arterial geometries such as the aortic arch. Using these patient specific geometries, we also investigate the importance of considering both the zero-pressure configuration and residual stresses on estimated material parameters and the stress distribution throughout the bifurcation, thereby providing insights into the use of stress as a possible plaque/vessel rupture vulnerability metric.

## 2. Methods

### 2.1 In -vivo MR Imaging Protocol

*In Vivo* MRI scans of the carotid arteries were obtained from healthy volunteers using a 3T whole body MRI scanner (Achieva, Phillips Medical Systems, Best, Netherlands) combined with an 8-channel dedicated bilateral carotid artery coil (Shanghai Chenguang Medical Technologies, Shanghai, China). The imaging parameters used for the creation of the geometries are stated in Table 2 with the field of view (FOV) centred on the carotid bifurcation after localization using the time of flight (TOF) sequence. For each volunteer, cuff diastolic and systolic pressures were recorded as shown in Table (1) and used as loading conditions in our finite element simulations. A triggering time was imposed so imaging could take place at both the diastolic and systolic phases of the cardiac cycle. This changed the overall scan time for each patient as it was dependent on their heart rate.

**Table 1:** Information acquired from volunteers

| Acquired Information | Volunteer #1 | Volunteer #2 | Volunteer #3 |
| --- | --- | --- | --- |
| Age (Years) | 25 | 30 | 42 |
| Diastolic Pressure (mmHg) | 75 | 68 | 82 |
| Systolic Pressure (mmHg) | 112 | 107 | 120 |

**Table 2:** Image acquisition parameters used for visualizing the vessel wall

| Acquisition Parameters | TOF | T1 | T2 | TSE |
| --- | --- | --- | --- | --- |
| No. of Slices | 48 | 8 | 8 | 8 |
| TE (ms) | 25 | 984 | 3000 | 2 R-R Intervals |
| TR (ms) | 3 | 11 | 38 | 38 |
| Resolution (mm) | 0.5 x 0.5 | 0.5 x 0.5 | 0.5 x 0.5 | 0.5 x 0.5 |
| Slice Thickness (mm) | 3 | 3 | 3 | 3 |
| No. of Echoes | 1 | 1 | 1 | 6 |
| Average Scan Time (mins) | 2:06 | 6:30 | 6:38 | 6:38 |

To test the impact of image resolution on our models, the field of view was translated by 1mm in the z-direction and then shifted 1mm again with acquisitions at both locations. This created a higher resolution with an “apparent” slice thickness of 1mm. To do this, the images were reallocated into the image stack, with a registration landmark set at the centre of the lumen ensuring that the images were correctly aligned so accurate segmentation could be performed.

### 2.2 Segmentation, Preparation and Hexahedral Meshing of Arterial Bifurcations

#### 2.2.1 Segmentation

After image acquisition, the DICOM stacks were input into Simpleware ScanIP (Synopsys, Inc., Mountain View, USA) for segmentation. The stack was first cropped to the region of interest and a mask was defined to perform the segmentation. Curves were delineated and arteries were then segmented manually from T2 weighted images of the carotid artery due to the high contrast of the vessel wall compared to the lumen and the surrounding tissue, see figure 1A. Each slice was analysed independently to ensure that the segmentation was as accurate as possible. This method accounted for the variable wall thickness of the vessel wall.

**Figure 1:**
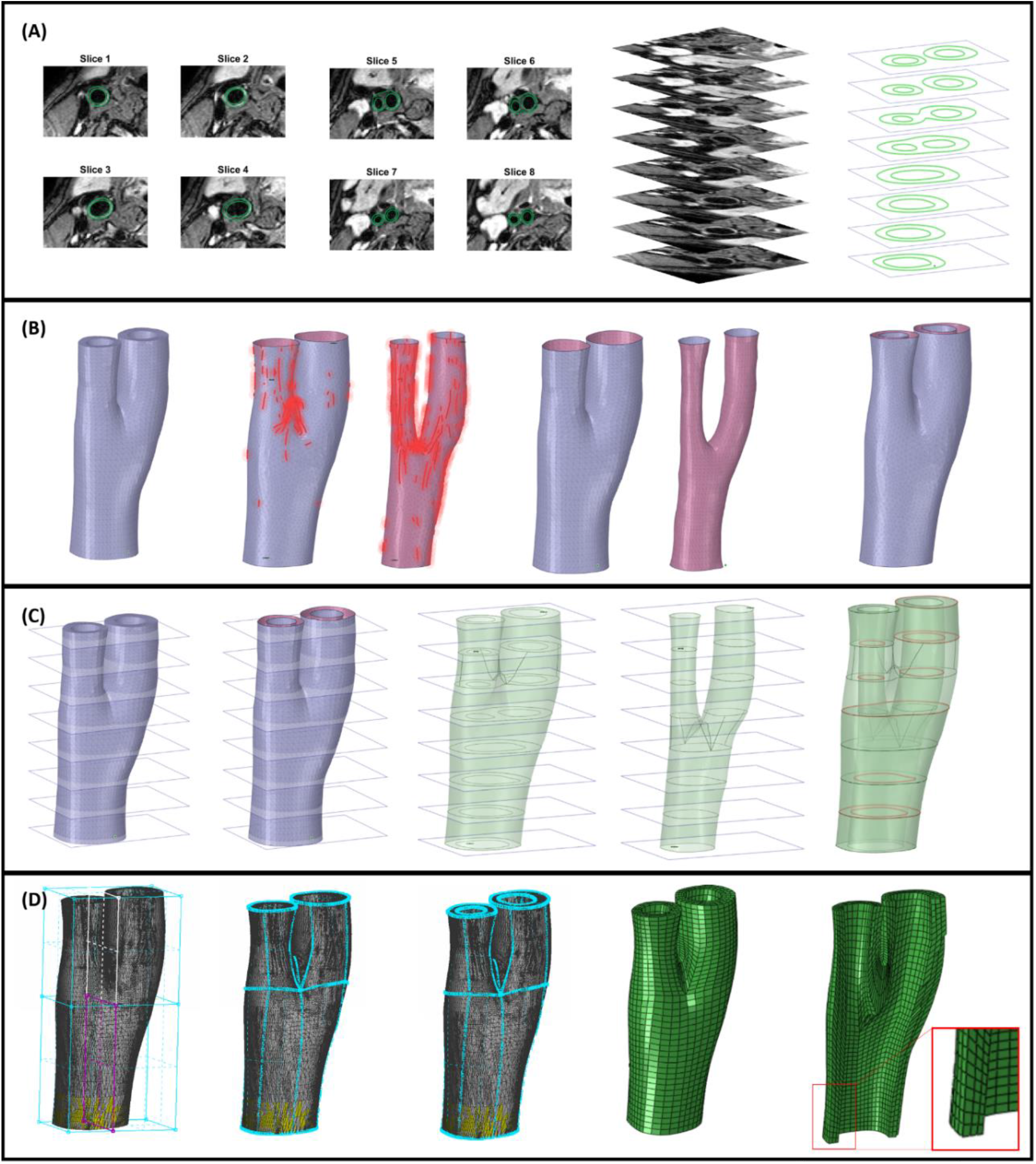
(A) Delineation of vessel from T2 weighted MRI images and creation of 3D stack (B) Initial model created – determining the location of sharp edges and smoothing the geometry (C) Surfaces of new smoothed surface are extracted and volume model is created for export into mesh pre-processor (D) Hexa-block tool used to define the geometry and create hexahedral finite element meshes of the bifurcation.

**Figure 2:**
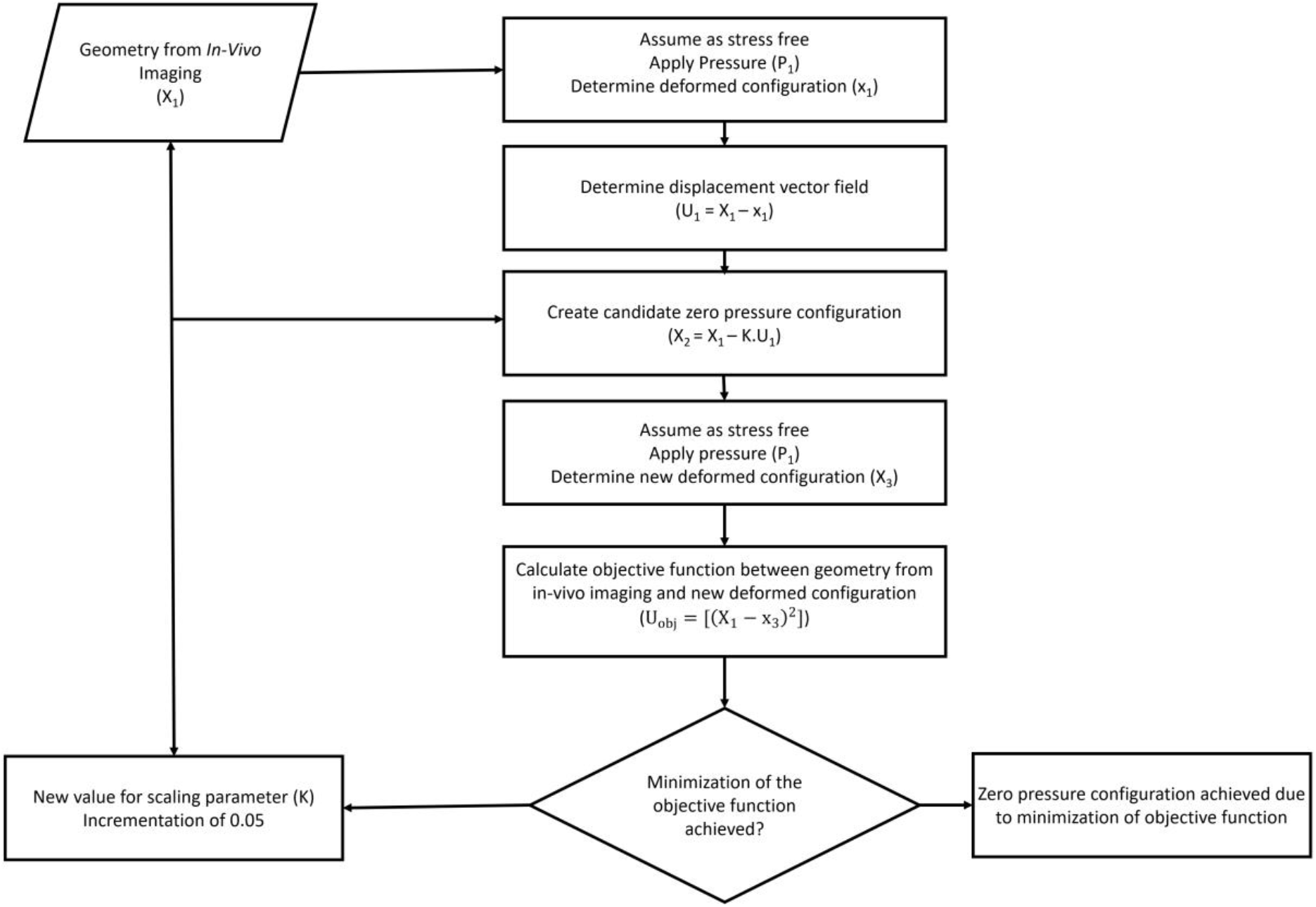
Algorithm workflow for extracting the zero-pressure configuration from a vessel geometry with known pressure conditions.

#### 2.2.2 Geometry Preparation

Once created, the segmented geometries needed to be smoothed further to avoid sharp edges in the reconstructed artery. This was especially important at the location of bifurcations, where an un-smoothed geometry could result in elements that cause not only numerical convergence issues but locations of high stress. Using ANSYS Spaceclaim (ANSYS Inc, USA), the geometry was imported and the STL facets were checked using the inspect function. Sharp edges, as shown in figure 1B, were firstly highlighted and subsequently smoothed using the fix sharps tool. After all sharp edges were removed, curves were extracted from the boundaries of the vessel wall by obtaining cross sections of the geometry using a series of parallel planes, see figure 1C. These curves were then connected to construct the inner and outer surfaces of the arterial wall. The geometry was then stitched together to ensure there were no gaps between surface interfaces in the next step. The blend function was then applied to connect the inner and outer vessel wall to ensure a connected geometry. These smoothed surfaces were then saved as STL meshes to export to ANSA pre-processor software.

#### 2.2.3 Hexahedral Meshing

After importing the new processed STL geometry, hexahedral meshing was performed using ANSA pre-processor software (v17.0, BETA CAE Systems, Thessaloniki, Greece). Using the *Hexa Block* module, the volume of the STL was defined initially in the form of a box, see figure 1D. This box was first split into four sections and the outside perimeters of the boxes were then moved to fit to the curvature of the geometry. Crosshatch faces, which are the intersecting planes between adjacent boxes, were then selected to separate the joined boxes in order to fit three independent boxes defining the three different sections of the carotid artery, more specifically, the common carotid, internal and external carotid. The perimeters of each defined box were then assigned to the outer wall of the geometry using the *project to surfaces* tool while the edges at proximal and distal were assigned using the *project to edges* tool. To include the inner wall, the O-Grid function was used, whereby inner perimeters were created and assigned to the interior wall, again using the *project to surfaces* and *project to edges* tools. To complete meshing, the connecting hatches in the lumen of the model were removed leaving just the assigned boxes to the vessel wall. This step is not necessary if fluid-based models are desired. Finally, the *Pure Hexa* module was applied to automatically generate a structured hexahedral mesh of the bifurcation with the desired mesh density. This mesh density could be set by setting the desired number of elements on the perimeters of the assigned boxes.

##### Constitutive Equation

For implementation in all models, the fibre reinforced HGO hyperelastic material model was used [64]. This model assumes that the tissue is composed of a matrix material that is embedded with two families of fibres, each has a preferred fibre direction. Mathematically, the model can be expressed by the strain energy function **W**:

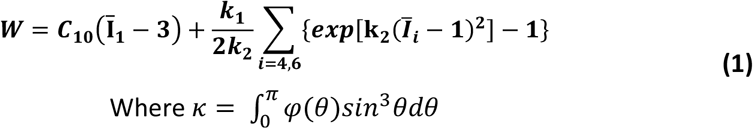

Where *C*_10_ is used to describe the isotropic matrix material along with the deviatoric strain invariant *Ī*_1_. k_1_is a positive material parameter with the units of kPa while k_2_ is a dimensionless parameter. The deviatoric strain invariant *Ī*_t_ is used to characterize each fibre family and κ is used as a dispersion parameter describing the fibre distribution, whereby κ = 0 describes high alignment while κ = 0.33 means the fibres are isotopically distributed. Θ is defined as the angle between the fibre direction and the circumferential axis. Five material parameters are therefore required to be input for the models and the values used are those in Balzani et al, 2012 and are reported in Table 3 [65].

**Table 3:** Material parameters used in computational models [65]

| $C_{10}$ (kPa) | $k_1$ (kPa) | $k_2$ | $\kappa$ | $\theta$ (deg) |
| --- | --- | --- | --- | --- |
| 6.56 | 1482 | 561 | 0.16 | 37 |

### 2.3 Zero Pressure Configuration

The method for estimation of the zero pressure configuration is detailed in the workflow illustrated in figure 2, and is adapted from Raghavan et al, 2006 [32]. Implemented in MATLAB (MATLAB 2019a, The MathWorks, Inc., Natick, Massachusetts, United States), X is the N x 3 array that describes the original undeformed nodal coordinate positions (x, y and z) and x is denoted to represent the deformed configuration after pressure loading. Element connectivity is kept constant throughout ensuring that no errors were incurred between initial and deformed configurations. Illustrated by the workflow in figure 9, the *in-vivo* geometry was extracted using the protocols described in Section 1. This initial *in-vivo* geometry **(*X***_**1**_)is initially assumed to be stress free and loaded with *in-vivo* pressure conditions **(*P***_**1**_). To create our candidate zero pressure configuration, the displacement vector field **(*U***_**1**_)was calculated. This displacement field vector is a subtraction of the initial *in-vivo* geometry and the deformed configuration **(*x***_**1**_).

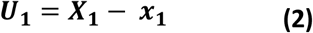

Once calculated, the displacement field is scaled using a parameter **K** and subtracted from the *in-vivo* geometry to obtain the first estimate of the zero-pressure configuration **(X**_**2**_).

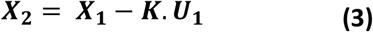

Here, it is again assumed that this first estimate of the zero-pressure configuration is stress free. The same pressure condition ***(P***_**1**_)is imposed as in the previous step to obtain a new deformed configuration **(*x***_**3**_).

Theoretically, this new deformed configuration **(*x***_**3**_) should be equal to the in-vivo geometry **(*X***_**1**_)from our initial step. However, due to the mechanical properties, the deformation of the geometry will be different. Therefore, to improve accuracy of the geometry estimated, an objective function of the displacement **(*U***_***obj***_) between the initial in-vivo geometry **(*X***_**1**_)and the deformed zero-pressure **(*x***_**3**_) configuration is implemented.

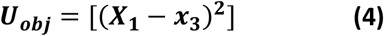

***U***_***obj***_ is then minimised until it reaches a set tolerance of less than 0.05 mm. If the geometry is not deemed optimised after a pre-set criterion of 50 iterations, a new value of K is implemented by incrementing the previous value by 0.05.

### 2.4 Residual Stress Implementation

For the entire mathematical derivation, the reader is directed to Schroder et al, 2014 [49]. Details included here are a summary of the algorithm implemented in that publication. For implementation into our models, we assumed that arterial tissue is an incompressible material. For fibre reinforced tissues such as arteries, we additively decomposed the Cauchy stress tensor ***σ*** into deviatoric ground stresses ***σ***^∗^ and reaction stresses ***σ***^***reaction***^, which represent the influence of the fibre stress state.

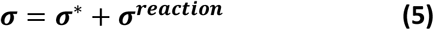

It is then assumed that this reaction stress is composed of the hydrostatic pressure imposed, the fibre tensions ***T***_**1**_ and ***T***_**2**_ and the structural tensors ***m***_**1**_ and ***m***_**2**_ which incorporate the preferred fibre directions ***a***_**1**_ and ***a***_**2**_.

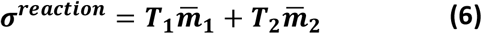

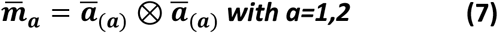

For the fibre tensions, ***T***_**1**_ and ***T***_**2**_, these can be obtained by using equation 6 and 7 respectively. Here the abbreviations 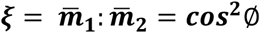, where *∅* is the inclination angle between the two fibre directions.

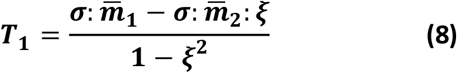

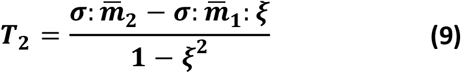

For physiological loading conditions, local volume averages of the fibre stresses are calculated for each decomposed volume segment ***v***^***mat***^

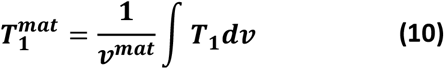

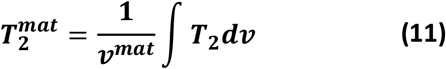

With the difference between this mean value and the fibre stresses giving equations 12 and 13.

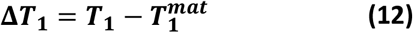

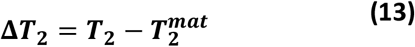

Where we now estimate the residual stresses to be:

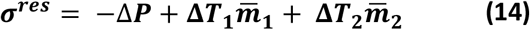

By which 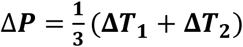

Now that we have calculated our residual stress tensor ***σ***^***res***^, we can subtract this to get our inclusion of our residual stresses in our Cauchy stress tensor by:

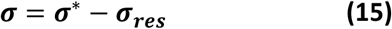

The method implemented in this study follows the algorithmic approach detailed in Schroder et al, 2014. Extended to 3D models, the residual stresses are incorporated by including a tensional fibre stress to homogenize the stress gradient throughout the vessel wall. Implemented via Abaqus UMAT and URDFIL subroutines connected by common blocks to allow the exchange of variables such as section volume, the algorithm is as follows

a. Segment the arterial wall into volume sections (see figure 3). Done in ANSA pre-processor software.

**Figure 3:**
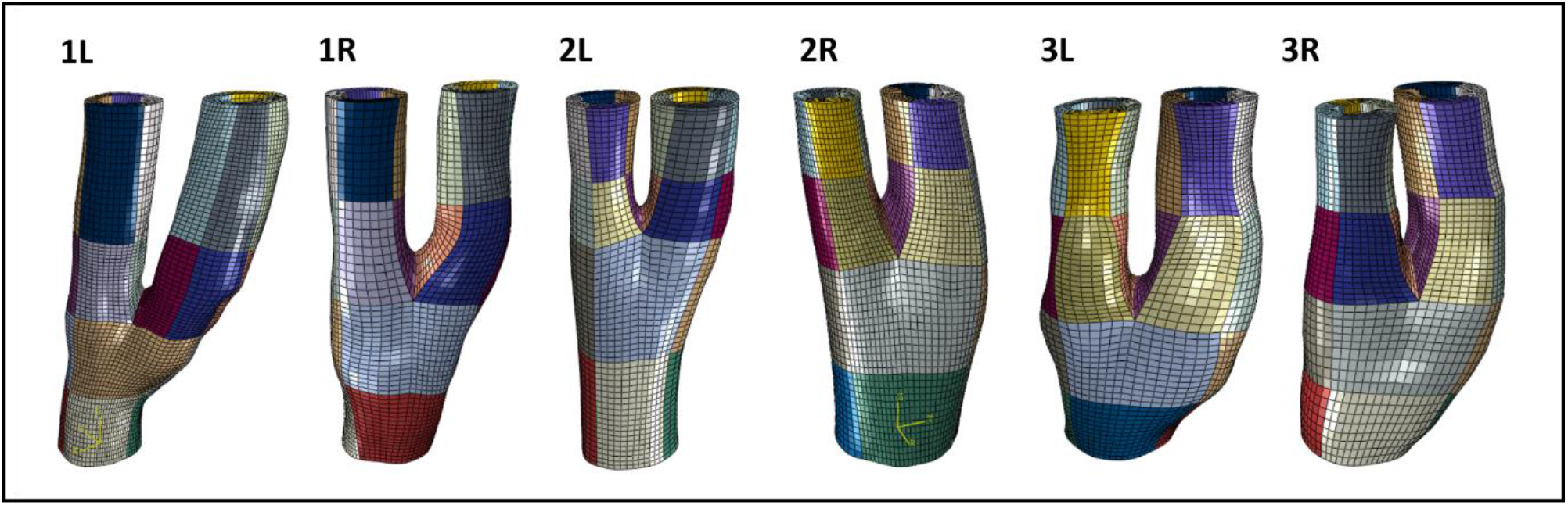
Defining volume sections throughout right bifurcation models. The material is defined in each section and volumes are calculated. L denotes that the bifurcation is on the left side while R denotes that the bifurcation is on the right side.
b. Define a suitable stress measure (tensional fibre stress).
c. Compute the local volume averages of the stress measure and the deviation of local and averaged stress measure in a section of interest.
d. Define the amount of residual stresses and apply a proportionate value to the equilibrium stress state (state at which geometry is loaded and not considering the residual stresses).
e. Iterate until convergence of the stress measure is observed.

Like Schroder et al, 2014, steps (c-e) are denoted as a “smoothing loop”; by which the smoothing loop is repeated until convergence of the solution is reached when no significant changes in the residual stresses are observed.

### 2.5 Inverse Material Parameter Estimation Implementation

The method of parameter estimation in this study is adapted from the implementation in Liu et al, 2017 [60]. The following statements are made to provide clarity in the method implemented:

a. *In-vivo* loaded geometries are extracted from two known phases in the cardiac cycle (.e.g., Diastole and Systole)
b. Finite element meshes at the two phases are constructed to ensure they have mesh correspondence, i.e., consistent number of elements and connectivity. This ensures that the displacement field can be obtained correctly
c. Thickness of the vessel wall can be directly inferred from MR Images.
d. Assuming that the vessel is statically determinant, assigning linear elastic properties (E = 2 × 10^4^ GPa and v = 0.49 [60]) will give the “almost true” stress state of the vessel, within 10% of the actual true stress experienced by the vessel

The inverse finite element algorithm workflow for material parameter estimation is illustrated by the schematic in figure 4 and is performed in Isight (Dassault Systemes Simulia corporations, Velizy-Villacoublay, France). This approach leverages that the “almost true” stress field of the vessel wall can be approximately determined using linear elastic properties.

**Figure 4:**
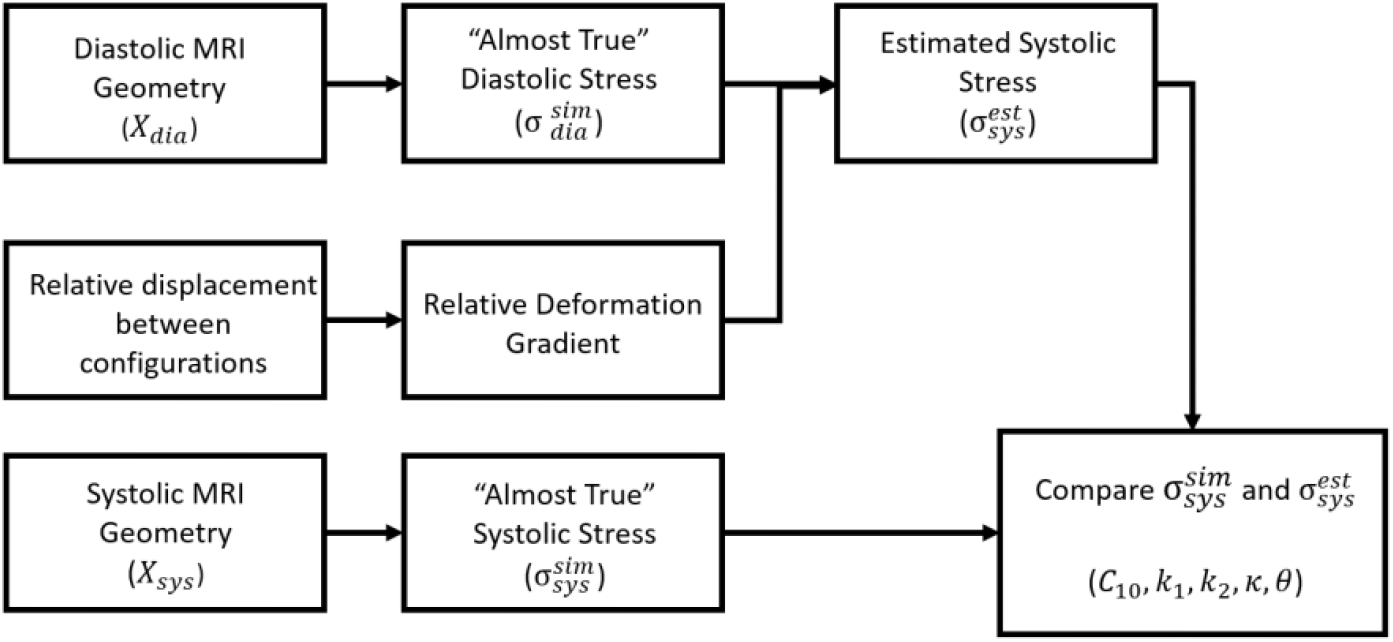
Algorithm workflow of the implemented inverse FE algorithm for material calibration of the carotid artery in Isight.

Different parameterized Abaqus input files have been designated accounting for the diastolic, systolic and zero pressure configuration cases. The input files are imported into the Abaqus component of Isight with the material parameters set as optimization variables. The optimization process is then formulated as follows: the objective is to find a set of constitutive parameters (*C*_10_, *k*_1_, *k*_2_, *κ, θ*), for the element type C3D8H in Abaqus. The square stress at all integration points are calculated and summed together as the value of the objective function as stated in the equation:

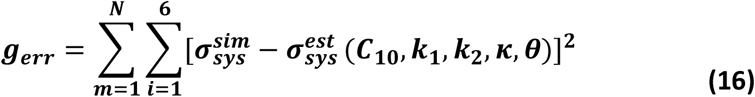

Where 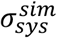 is the “almost true” stress state and 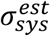 is the estimated stress state with assigned material parameters. N is the number of elements used in the optimization. Upper and lower bounds stated in table 4 are set for all parameters.

**Table 4:** Lower and higher bounds set for material parameter estimations. Material parameters used here are extracted from Balzani et al, 2012 [65].

| Parameters | $C_{10}$ (kPa) | $k_1$ (kPa) | $k_2$ | $\kappa$ | $\theta$ (deg) |
| --- | --- | --- | --- | --- | --- |
| **Lower Bound** | 0 | 0 | 0 | 0 | 0 |
| **Initial Input Parameters** | 6.56 | 1482 | 561 | 0.16 | 37 |
| **Higher Bound** | 20 | 2000 | 1000 | 0.33 | 45 |

**Table 5:**
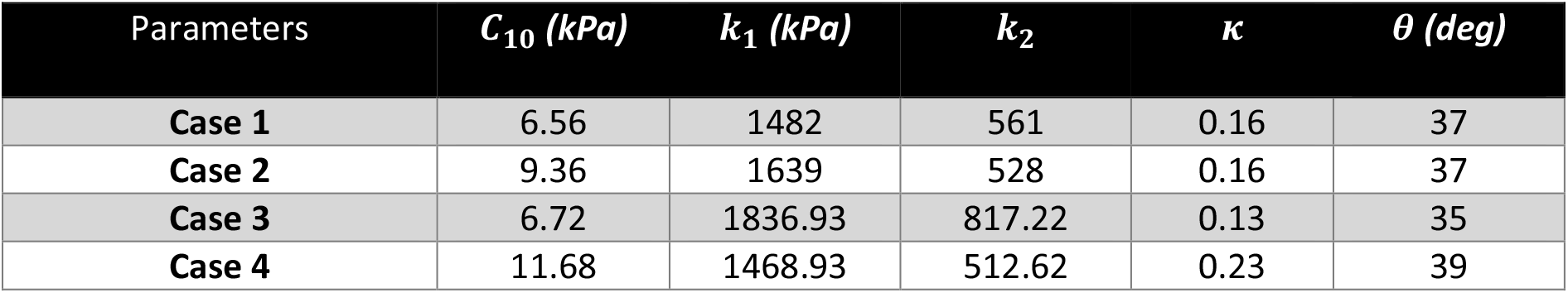
Implemented material properties in each case for one geometry

| Parameters | $C_{10}$ (kPa) | $k_1$ (kPa) | $k_2$ | $\kappa$ | $\theta$ (deg) |
| --- | --- | --- | --- | --- | --- |
| Case 1 | 6.56 | 1482 | 561 | 0.16 | 37 |
| Case 2 | 9.36 | 1639 | 528 | 0.16 | 37 |
| Case 3 | 6.72 | 1836.93 | 817.22 | 0.13 | 35 |
| Case 4 | 11.68 | 1468.93 | 512.62 | 0.23 | 39 |

To test the effect of including the zero-pressure configuration and residual stress on estimated material parameters and the overall stress distribution throughout the bifurcation, a number of cases are considered:

**Case 1:** Models extracted at diastole are simulated with literature material parameters
**Case 2:** Models are simulated with estimated material parameters from diastole to systole
**Case 3:** Models start in the zero-pressure configuration with estimated parameters going from this zero-pressure configuration to diastole and systole
**Case 4:** Residual stress is included at zero-pressure configuration and simulated with estimated material parameters from the zero-pressure configuration to diastole and systole.

To further investigate the effect of including the zero-pressure configuration and residual stress on the stress strain response, a single element attributed with estimated material parameters is simulated in uniaxial tension. Furthermore, percentage volume plots are extracted from the bifurcation models to observe the effect these implemented methods have on the stress distribution

## 3. Results

### 3.1 Effect of Image resolution on Stress Calculations

To demonstrate the sensitivity of our models to stresses calculated, it is important to do a mesh convergence analysis. This is to ensure that the most accurate stress estimation can be calculated in the least amount of computational time without compromising the accuracy of the result. In this mesh convergence analysis, the location of highest stress was the location of interest and like the literature and from these biomechanical models, highest stresses were observed at the location of the apex of the bifurcation.

For robust mesh convergence analysis, it would be logical to argue that for models of similar geometry, then the same level of mesh refinement would be suitable to obtain the same level of accuracy. Therefore 250,000 elements are deemed enough for our analysis as values do not change by more than 5% of the previous value, see figure 5. Interestingly, it is observed that convergence of stress results is independent on the model resolution. For our low-resolution model, the convergence is observed at a stress value of 286 kPa while for the high-resolution model the convergence is observed at stress values of 145 kPa. The sensitivity difference between these two observed values is 49%, which is a significant increase and would lead to an overestimation of the stresses at the apex location.

**Figure 5:**
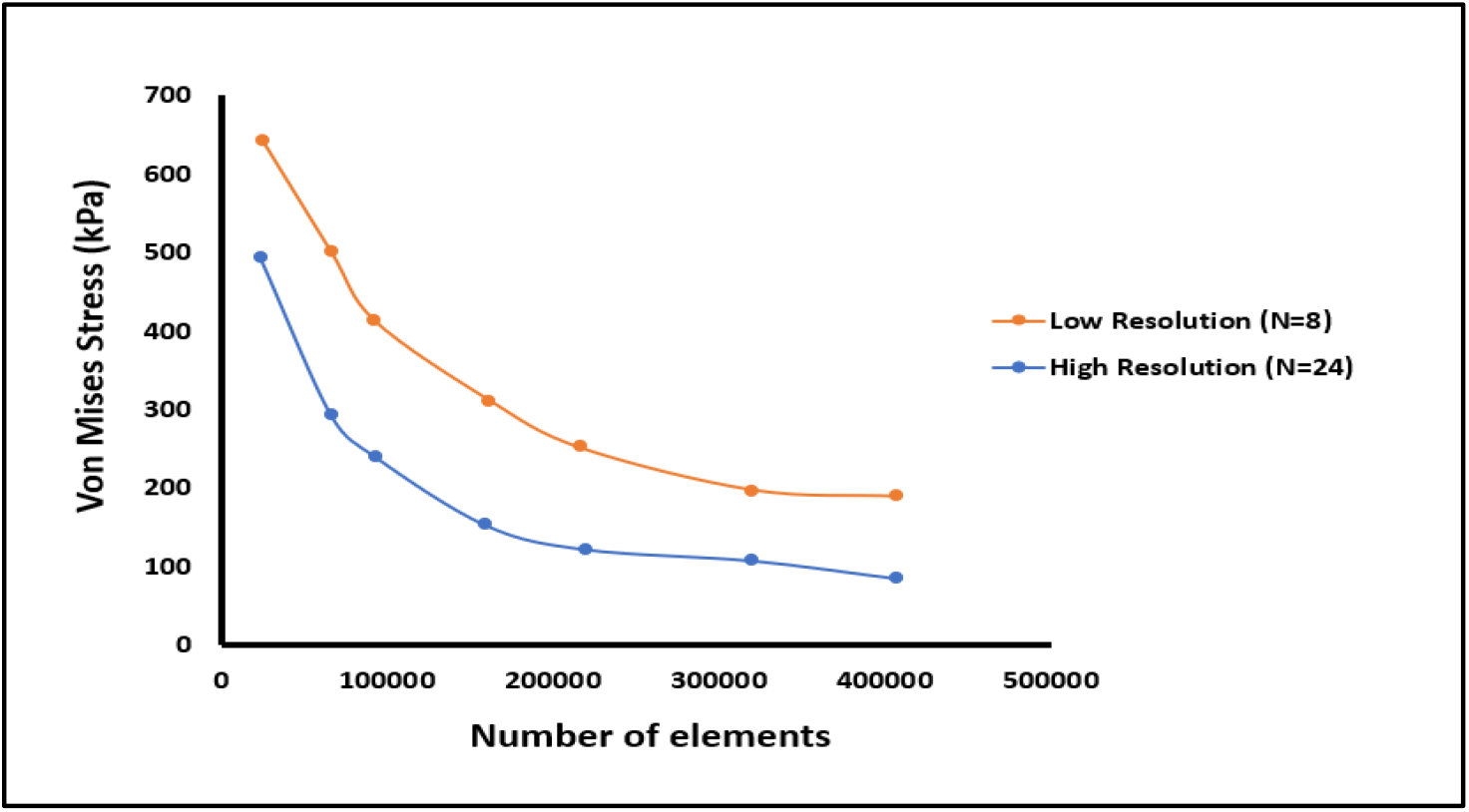
Mesh convergence results for bifurcation geometry 2L at two different resolutions (a) Low resolution model where Number of slices (N) = 8 (b) High resolution model where Number of slices (N) = 24

### 3.2 Optimization of the Zero Pressure Configuration

#### 3.2.1 Cylinder Model

For our estimation of the zero-pressure configuration, an idealized cylinder model is first used to test the robustness of the implemented algorithm. For our idealized model, the wall thickness was set to be 0.75mm with an inner diameter of 2mm. An internal pressure was also assigned to be 16kPa. As demonstrated by figure 6A, the estimated zero pressure configuration comprises of a lower luminal radius and larger wall thickness when compared to the initial loaded configuration. This is expected and follows results demonstrated in Raghavan et al, 2006 [30].

**Figure 6:**
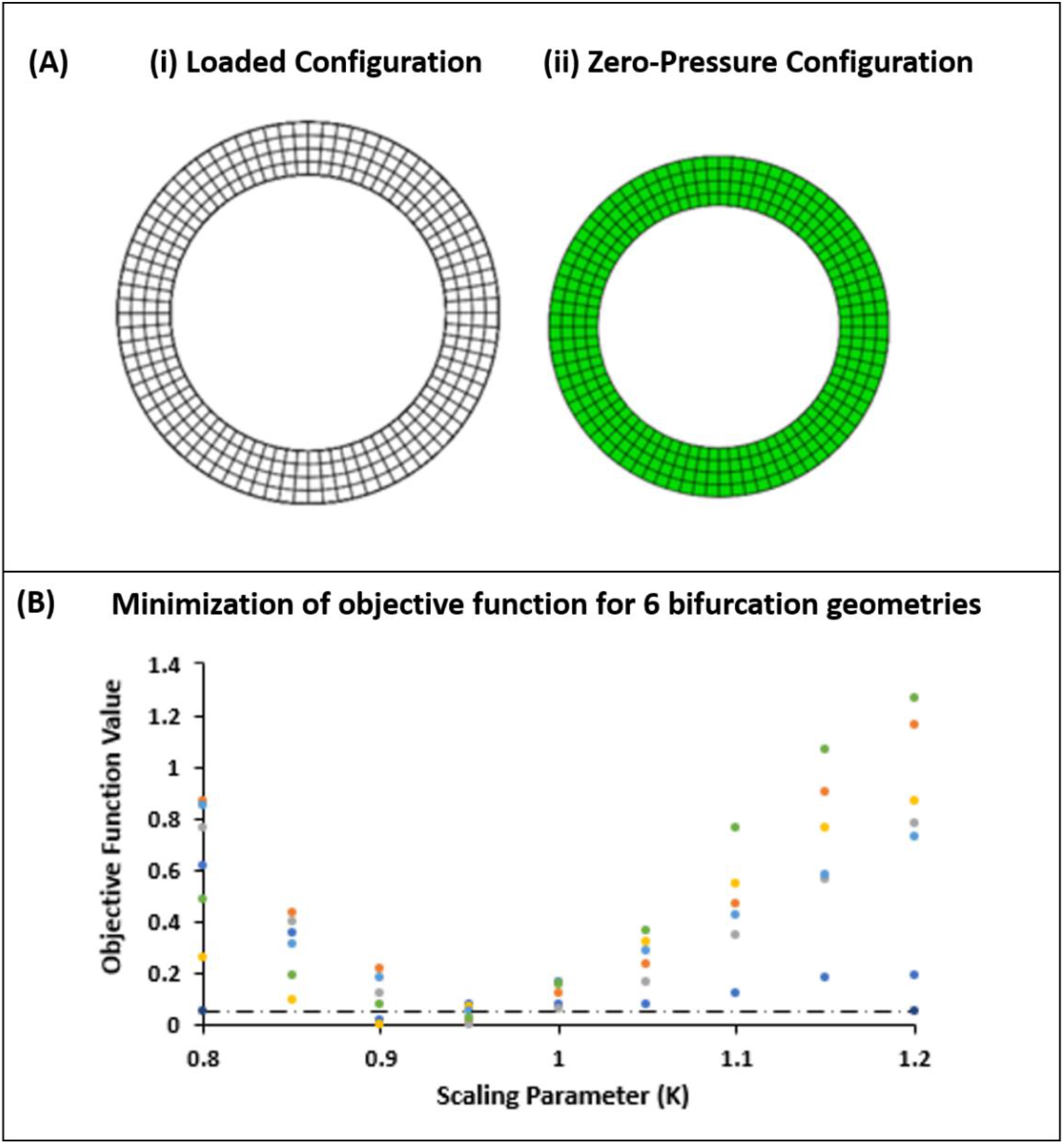
(A) Optimization of the zero-pressure configuration for an idealized cylinder model. The wireframe shown in (i) depicts the initial geometry while the green geometry shown in (ii) shows the estimated zero pressure configuration. (B) Optimization of the zero-pressure configuration for patient specific carotid bifurcation models

#### 3.2.2 Bifurcation Models

As shown by figure 6B, the algorithm shows convergence of the used scaling parameter K to be in between 0.9 and 1 for all 6 bifurcation models, which is similar to results obtained in Raghavan et al, 2006 for an aneurysmal model [30]. After finding the optimal zero-pressure configuration, the wall thickness of the bifurcation models increased with a decrease in luminal radius. It is important to note that lower increments <0.05 increases the accuracy of the estimated zero pressure configuration. However, due to a new simulation being needed after each iteration, this would increase the computational time needed.

### 3.3 Optimization of residual stress inclusion

To validate the accuracy of the implemented algorithm, a simplified cylindrical model was used. The reason for this is to have an approximate value of stress to optimize towards otherwise a complete smoothing of the gradient would be achieved. Using classical mechanics, it was possible to calculate the expected stress manually before our simulation to validate the calculated stresses observed in our model. By assuming that the model is a thin-walled cylinder, the circumferential stress can be calculated by:

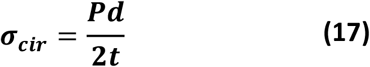

Where P is the pressure imposed, d is the inner diameter of the model and t is the wall thickness. For our model, the wall thickness was 0.75mm with an inner diameter of 2mm. Pressure imposed for the model was set to be 120mmHg which corresponds to approximately 16 kPa. The circumferential stress calculated for this cylindrical model was 21.3 kPa, see figure 7A

**Figure 7:**
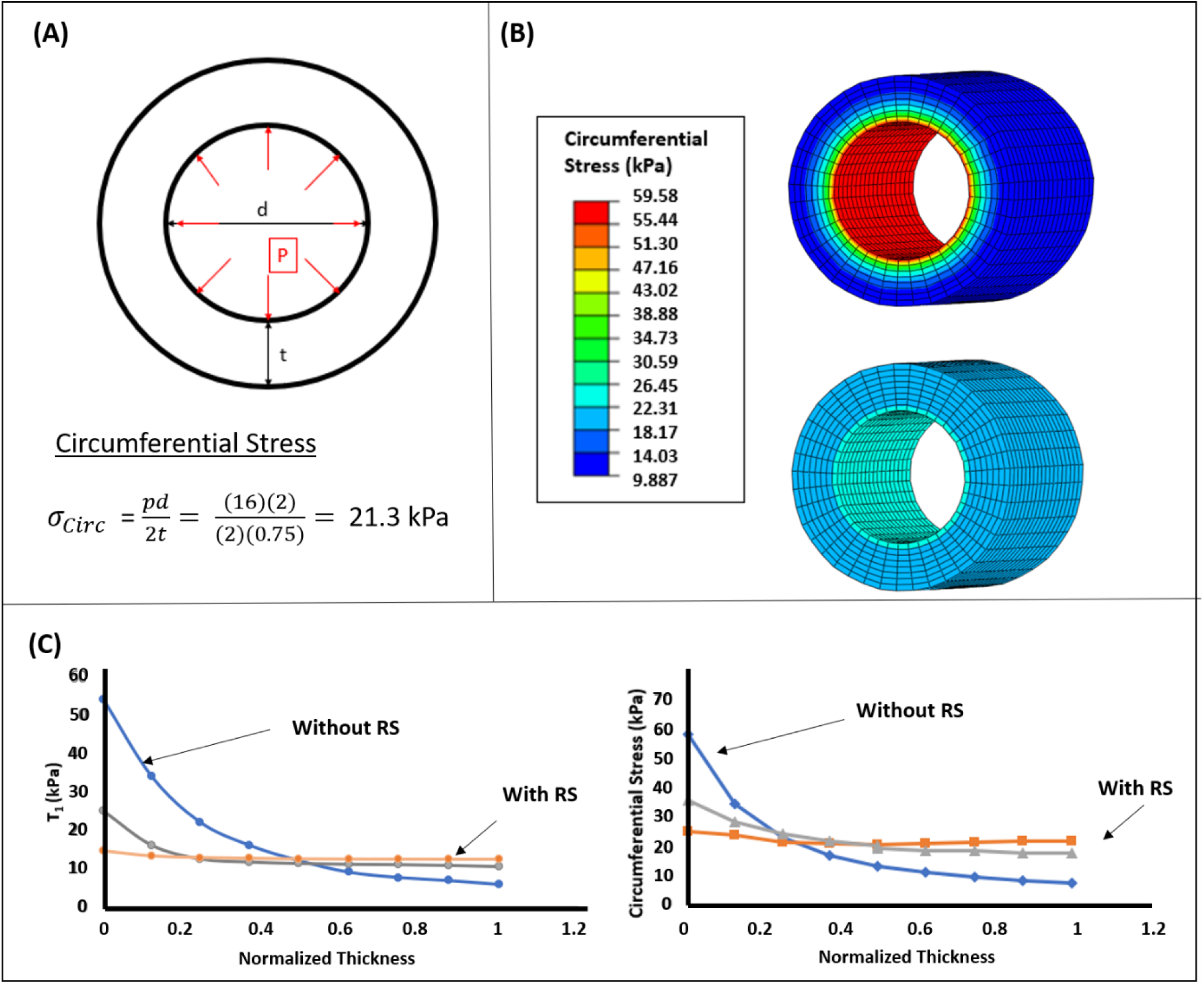
Inclusion of residual stresses in our cylindrical model (A) Theoretical calculation of the circumferential stress (B) Circumferential stress contour plots showing (i) without the inclusion of residual stress and (ii) with the inclusion of residual stress (C) (i) Fibre stress T_1_ (kPa) over the normalized radius (mm) after applying 0, 5 and 10 smoothing loops (ii) Impact of inclusion of residual stresses on the maximum principal stress (kPa) after applying 0, 5 and 10 smoothing loops

#### 3.3.1 Cylinder Model

Using the same dimensions as our theoretical model, both the tensional fibre stresses ***T***_**1**_, ***T***_**2**_ and maximum principal stress gradients decrease when imposing an increased number of smoothing loops. Residual stress was deemed to be incorporated into the model when the tensional fibre stress ***T***_**1**_ converged but preserved the small gradient of maximum principal stress from the inner to outer wall, as shown by figure 7C. Tensional fibre stress ***T***_**2**_ is equal to ***T***_**1**_. 10 smoothing loops was determined to be enough to incorporate residual stresses into our cylindrical model as the tensional fibre stress converged when the tensional fibre stress from inner to outer wall was within 10%. For our cylindrical models, the maximum principal stress from outer to inner wall goes from 25 kPa to 21kPa with the inclusion of residual stresses as illustrated in figure 7B. This is in the range of our theoretical calculation that the vessel wall experiences a stress of approximately 21.3kPa.

#### 3.3.2 Bifurcation Model

To determine the optimum number of smoothing loops needed for our bifurcation models, the process was extended to include additional smoothing loops until values of less than 5 kPa were seen in the tensional fibre convergence. The location of interest is the bifurcation apex, where we see the largest gradient of stress through the vessel wall.

Like the results obtained in our simplified cylinder model. Both the tensional fibre stress ***T***_**1**_ and maximum principal stress gradients decrease when imposing an increased number of smoothing loops, see figure 8C and figure 8D. The gradient of stress seen initially is considerably larger than our cylindrical model and therefore requires more smoothing loops for our tensional fibre stress to converge to our pre-set criterion. Furthermore, a reduced gradient of stress is preserved from inner to outer wall, demonstrating we have included the residual stress in our models. Again theoretically, the predicted stress at locations such as the common, internal and external carotid are similar to the predicated values after the inclusion of the residual stress, see figure 8A and 8B.

**Figure 8:**
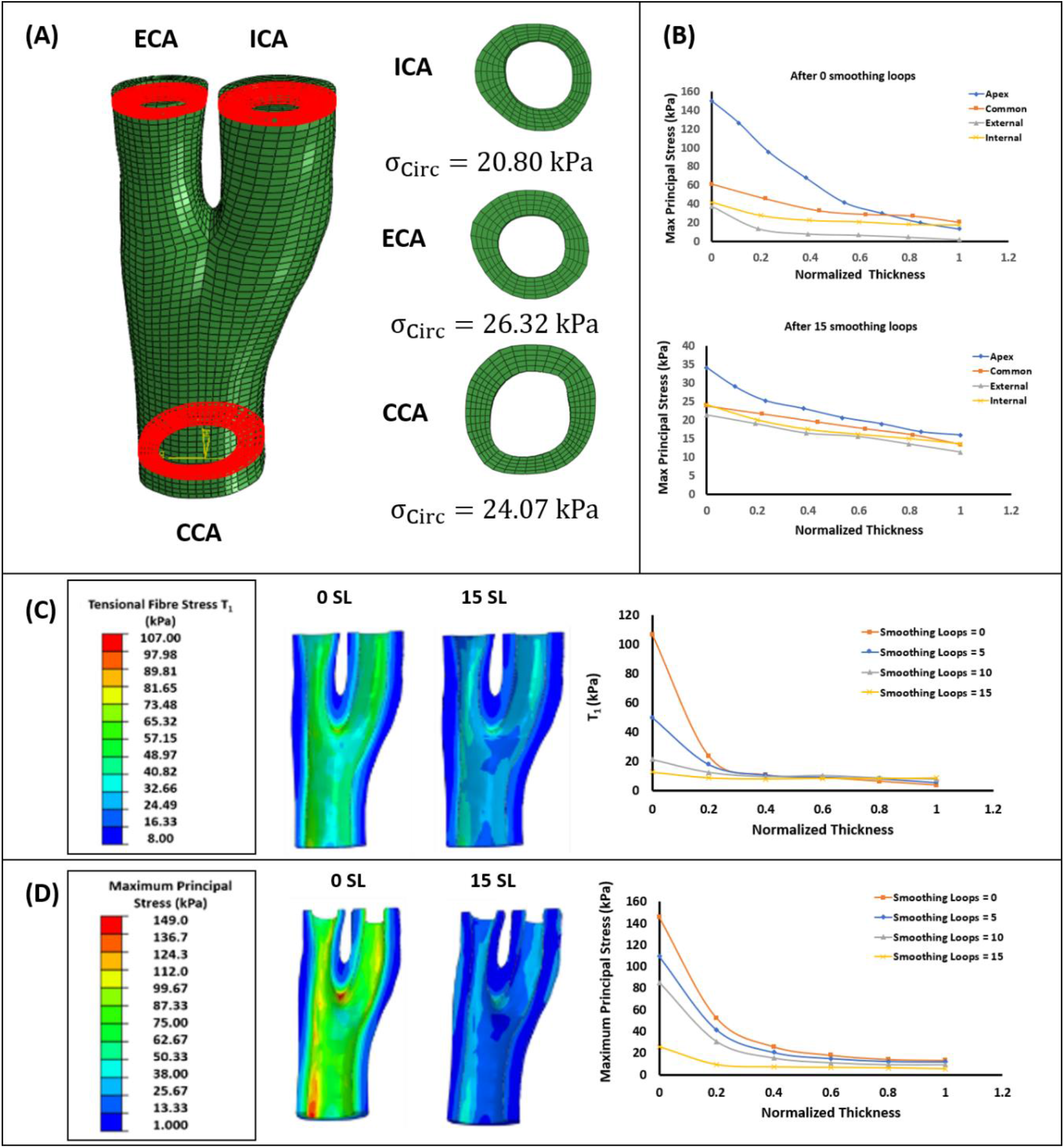
(A) Locations across the bifurcation for theoretical approximation of the circumferential stress (B) Inclusion of the residual stress at the common, external and internal carotid (C) Inclusion of residual stresses in the bifurcation model looking at the Fibre stress T_1_ (kPa) over the normalized radius (mm) after applying 0, 5, 10 and 15 smoothing loops (D) Impact of inclusion of residual stresses on the maximum principal stress (kPa) after applying 0, 5, 10 and 15 smoothing loops

### 3.4 Estimation of Material Parameters and Calculation of the Stress Distribution

Presented in Figure 9 are the results obtained after simulating each case for one geometry. Six geometries were simulated in total and the reader is directed to the supplementary documentation provided for the simulation results on the other 5 geometries.

**Figure 9:**
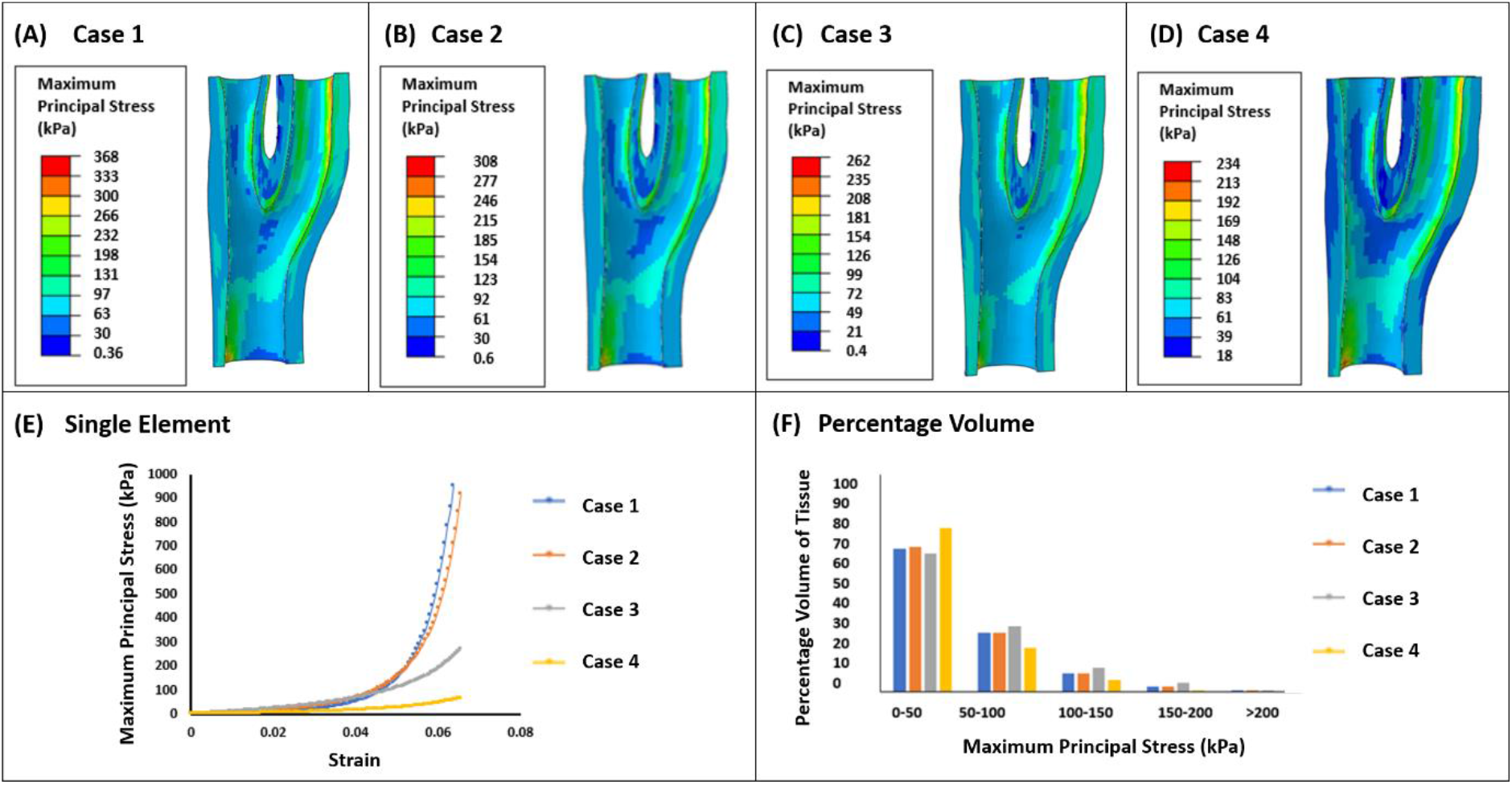
(A) Simulated stress result for case 1 (B) Simulated stress result for case 2 (C) Simulated stress result for case 3 (D) Simulated stress result for case 4 (E) Single element test of estimated material parameters under uniaxial tension (F) Percentage of volume graph for the 4 cases showing the stress distribution throughout the vessel wall

Figure 9 (A-D) demonstrates that the peak stress values are different in all cases considered, whereby the stress is highest with material parameters taken from literature (case 1) and the lowest when we include both the zero-pressure configuration and residual stress (case 4). Furthermore, this is observed in all geometries, see supplementary documentation. In case 2 material parameter estimations, the geometry taken from diastole is attributed material parameters from case 1 that need to be optimized and is loaded to systole. It is observed that due to the small deformation from diastole to systole that the most significant parameter change comes from *C*_10_, which describes the isotropic ground matrix of the tissue and is responsible for its initial response. In case 3 the material parameter estimations are determined when the zero-pressure configuration of the geometry is incorporated and then loaded to both diastole and systole. Due to the greater deformation of the geometry to the desired phases, the material parameters now start to alter more significantly. In this estimation, the parameters *C*_10_, k_1_ and k_2_ differ significantly from the original input parameters. *κ* and θ alter slightly but not significantly from the original input parameters. In case 4 material parameter estimations, where we included the zero-pressure configuration and residual stress it is observed that loading this to both diastole and systole, not only do they alter *C*_10_, k_1_, and k_2_ but also the dispersion parameter *κ*. In case 4, due to the significant change in *κ* the fibres become more dispersed when residual stress is included, decreasing the peak stress values observed in all regions of the bifurcation. Our single element results shown in Figure 9E, highlights the effects of the estimated material parameters with the inclusion of the zero-pressure configuration and residual stress. In all cases, the parameters taken directly from literature show a stiffer response then what is seen when we have optimized patient specific parameters, whereby it is observed that the inclusion of the zero pressure configuration and residual stress has a distinctly less stiff response. Lastly, the decrease in peak stress values and the stress distribution is further validated by the percentage volume plots shown in Figure 9F. It is observed that there is an increase in the percentage volume of the lower stress value band (0-50kPa) when the zero-pressure configuration and residual stress is included in the model (Case 4), depicting a less stiff response and lower peak stress values when compared to literature parameters (Case 1).

## 4. Discussion

### 4.1 Impact of Axial Resolution on Geometry stress calculation

As observed in our section 3.1 results, the high-resolution model with an “apparent” image slice thickness of 1mm reduced our calculated stress at the apex of the bifurcation models. This agrees with previous research undertaken by Nieuwstadt et al, 2014, whereby on a simulated MRI dataset they determined that lower stresses are predicted with higher resolution models [66]. To the authors knowledge, this investigation is the first to look at whether this also holds true for *in-vivo* MRI datasets. In terms of our created geometries, convergence of stress values at the bifurcation apex was observed at a mesh density of approximately 250,000 elements. Convergence was deemed to have been achieved when the stress values did not differ by more than 5% of the previous value. Interestingly, it was observed that for both the higher and lower resolution models, convergence was observed at different stress values. This is due to one geometry being created with aa higher resolution than the other. A limitation that should be considered however, is scanning duration as acquiring a high resolution may require longs scans and motion may cause reduced image quality over long scan times.

Overall, a robust technique for creating carotid bifurcation geometries from MRI has been established. From segmentation, preparation and meshing, these techniques can be translated to other arterial geometries, if required. The sensitivity of stresses calculated is dependent on the resolution of the acquisition that is used; therefore, a higher resolution scan is desirable.

### 4.2 Inclusion of the zero-pressure configuration

Using the method presented in section 2.3, the zero-pressure configuration can be extracted for idealized models and arterial bifurcations. For our idealized cylinder model, the robustness of the algorithm was demonstrated by recovering the estimated zero pressure configuration after inputting a known pressure state. The scaling parameter K was optimized at 0.90 to 1 in the idealized case. For our bifurcation models, the algorithm successfully estimated the zero-pressure configuration from geometries at a known phase in the cardiac cycle. Due to the geometry not being idealized, the scaling parameter needed to be optimized to determine the optimum incrementation to be used for robust estimation. This was found to be in the region of 0.9 and 0.95 for all geometries tested here, see figure 6. It was determined that an increment of 0.05 for new guesses of the zero pressure geometry delivered sufficient accuracy and proved to be most time effective for our models and is also the increment implemented in Raghavan et al, 2006 [32]. If stress is to be used as a clinical vulnerability indicator the zero-pressure configuration must be included for simulation of the arterial bifurcation.

### 4.3 Inclusion of Residual Stresses

The algorithm presented shows that we have homogenized the gradient of stress in the radial direction to represent the presence of residual stress in both our idealized cylinder model and our bifurcation models, see figure 7 and 8. Our idealized cylinder model shows the robustness of this method and can be compared to the 2D simulations in Schroder et al, 2014, where the gradient of stress in homogenized in the radial direction to a reasonable degree after 10 smoothing loops. The circumferential stress values observed through the wall thickness also falls in the range of the theoretically calculated value of 21.3kPa, further validating its implementation. The method presented here extends the method demonstrated in Schroder et al, 2014 to three-dimensional patient specific models of the carotid bifurcation. Including residual stress in our patient specific geometry again homogenizes the gradient of stress observed throughout the bifurcation but does not completely remove it. Particularly at the bifurcation apex, where we see the highest change in peak stress values when residual stress is included. The preservation of this gradient through the thickness is important and observed in the work of Delfino et al, 1997, Raghavan et al, 2003 and Alustrue et al, 2007 to name a few. It is important to remember that the residual stress inclusion shown in those studies used the opening angle approach, something that was not used in this implementation. The reason for this is because the opening angle varies through the bifurcation and cannot be quantified *in-vivo*. Therefore, using our approach, we neglect the opening angle, but show we can incorporate the residual stress of the vessel in an accurate manner.

### 4.4 Effect of implemented methods on estimated material parameters and stress distributions throughout the carotid bifurcation

From our results, the estimated material parameters alter with each geometry even without both the zero-pressure configuration and residual stress. This is due solely to the fact that the geometries are all different; this demonstrates the need to have patient specific material parameters. Looking at each geometry individually, it is observed that from diastole to systole (Case 2) that changes in the material parameters are mainly governed by the *C*_10_ parameter. When the zero-pressure configuration is included in our estimations, the geometry has to undergo further deformation to reach the diastolic and systolic phases, which effects the overall stress that is calculated. The introduction of the zero-pressure configuration also shows larger changes in the k_1_, k_2_ parameters along with an observed change in *C*_10_. It can be stated with the inclusion of the zero-pressure configuration that the contribution of the collagen fibres in the stress distribution are now being observed. Lastly, the inclusion of the zero-pressure configuration and residual stress alter the *C*_10_, k_1_, k_2_ and the dispersion parameter κ. As we know from the literature and our results, the presence of residual stress throughout the arterial wall homogenizes the gradient of stress through the wall thickness. This homogenization effects the parameter estimation and requires the fibres to be more disperse, therefore decreasing the overall stress of the model. This decrease in the stress gradient from inner to outer wall is seen in all models that includes the residual stress of the vessel. It is also observed that the maximum principal stress for each model is overestimated when simulated with material parameters taken from the literature. The stress values decrease when the zero-pressure configuration and residual stresses are included. This is important as it demonstrates the need for tailoring the analysis to be patient specific and including the zero-pressure configuration and the residual stress when estimating material parameters. Overestimation of the stress with parameters from literature and exclusion of the zero-pressure configuration and residual stress leads to a mischaracterization of the vulnerability of the vessel if stress is used as a vulnerability measure.

## 5. Limitations and Future Work

The work presented here shows the impact of excluding the zero-pressure configuration and residual stresses in the estimation of material parameters and stresses at the carotid bifurcation. The first limitation of the study is that the carotid wall is assumed to be made of one material. This is not the case as arterial tissue is composed of three distinct layers, the tunica intima, the tunica media, and tunica adventitia, all of which independently have their own opening angle, and therefore residual stresses. The protocol for geometry segmentation and meshing has been extended to include the meshing of multiple layers and plaque components and will be our future work. The second limitation of the study is exclusion of diseased vessel bifurcations in the analysis of the effect of the zero-pressure configuration and residual stresses. Although the residual stress does not have a major influence in atherosclerotic plaque tissue, the exclusion of the residual stress from the vessel wall with plaque tissue significantly alters the calculated stress values. Therefore, for more accurate simulation we should take out the plaque, include the residual stress in the vessel wall and re-incorporate the plaque back into the model. Lastly, the angle of fibres does not show considerable change in all estimations undertaken here. This could be from the fact that different ranges of axial strain were not implemented in our material parameter estimation and kept constant for all models. It is expected that if higher levels of axial strain are imposed, the angle of fibres would alter and would start to play a more significant role in the stress distribution and peak stress values.

## Ethical Approval

Ethical approval for obtaining the plaques in this study was obtained from St. James Hospital ethical committee in compliance with the declaration of Helsinki.

## References

[1] WHO, “Global status report on noncommunicable diseases 2010,” World Heal. Organ., p. 176, 2011.

[2] C. D. Mathers and D. Loncar, “Projections of global mortality and burden of disease from 2002 to 2030.,” PLoS Med., vol. 3, no. 11, p. e442, 2006.

[3] E. J. Benjamin et al., Heart Disease and Stroke Statistics 2017 Update: A Report from the American Heart Association, vol. 135, no. 10. 2017.

[4] B. A. Wasserman, R. J. Wityk, H. H. Trout, and R. Virmani, “Low-grade carotid stenosis: Looking beyond the lumen with MRI,” Stroke, vol. 36, no. 11, pp. 2504–2513, 2005.

[5] T. Hatakeyama, H. Shigematsu, and T. Muto, “Risk factors for rupture of abdominal aortic aneurysm based on three-dimensional study,” J. Vasc. Surg., vol. 33, no. 3, pp. 453–461, 2001.

[6] K. Gaba, P. A. Ringleb, and A. Halliday, “Asymptomatic Carotid Stenosis: Intervention or Best Medical Therapy?,” Curr. Neurol. Neurosci. Rep., vol. 18, no. 11, pp. 1–9, 2018.

[7] C. Huang et al., “Ultrasound-Based Carotid Elastography for Detection of Vulnerable Atherosclerotic Plaques Validated by Magnetic Resonance Imaging,” Ultrasound Med. Biol., vol. 42, no. 2, pp. 365–377, 2016.

[8] M. F. Fillinger, M. L. Raghavan, S. P. Marra, J. L. Cronenwett, and F. E. Kennedy, “In vivo analysis of mechanical wall stress and abdominal aortic aneurysm rupture risk,” J. Vasc. Surg., vol. 36, no. 3, pp. 589–597, 2002.

[9] T. C. Gasser, M. Auer, F. Labruto, J. Swedenborg, and J. Roy, “Biomechanical Rupture Risk Assessment of Abdominal Aortic Aneurysms : Model Complexity versus Predictability of Finite Element Simulations,” Eur. J. Vasc. Endovasc. Surg., vol. 40, no. 2, pp. 176–185, 2010.

[10] D. Tang et al., “Local critical stress correlates better than global maximum stress with plaque morphological features linked to atherosclerotic plaque vulnerability: An in vivo multi-patient study,” Biomed. Eng. Online, vol. 8, pp. 1–9, 2009.

[11] Z. Teng et al., “3D critical plaque wall stress is a better predictor of carotid plaque rupture sites than flow shear stress: An in vivo MRI-based 3D FSI study,” J. Biomech. Eng., vol. 132, no. 3, pp. 1–9, 2010.

[12] D. Tang et al., “Sites of Rupture in Human Atherosclerotic Carotid Plaques Are Associated With High Structural Stresses,” Stroke, vol. 40, no. 10, pp. 3258–3263, 2009.

[13] Z. Teng et al., “In vivo MRI-based 3D mechanical stress-strain profiles of carotid plaques with juxtaluminal plaque haemorrhage: An exploratory study for the mechanism of subsequent cerebrovascular events,” Eur. J. Vasc. Endovasc. Surg., vol. 42, no. 4, pp. 427–433, 2011.

[14] M. Ghasemi, R. D. Johnston, and C. Lally, “Development of a Collagen Fibre Remodelling Rupture Risk Metric for Potentially Vulnerable Carotid Artery Atherosclerotic Plaques,” Front. Physiol., vol. 12, no. October, pp. 1–17, 2021.

[15] M. Ghasemi, D. R. Nolan, and C. Lally, “Assessment of mechanical indicators of carotid plaque vulnerability: Geometrical curvature metric, plaque stresses and damage in tissue fibres,” J. Mech. Behav. Biomed. Mater., vol. 103, no. November 2019, p. 103573, 2020.

[16] A. Creane, E. Maher, S. Sultan, N. Hynes, D. J. Kelly, and C. Lally, “A remodelling metric for angular fibre distributions and its application to diseased carotid bifurcations,” no. 2006, 2011.

[17] G. De Santis, M. De Beule, K. Van Canneyt, P. Segers, P. Verdonck, and B. Verhegghe, “Medical Engineering & Physics Full-hexahedral structured meshing for image-based computational vascular modeling,” Med. Eng. Phys., vol. 33, no. 10, pp. 1318–1325, 2011.

[18] L. Antiga, B. Ene-Iordache, L. Caverni, G. P. Cornalba, and A. Remuzzi, “Geometric reconstruction for computational mesh generation of arterial bifurcations from CT angiography,” Comput. Med. Imaging Graph., vol. 26, no. 4, pp. 227–235, 2002.

[19] Seung Lee et al., “Automated mesh generation of an arterial bifurcation based upon in vivo MR images,” pp. 719–722, 2002.

[20] G. De Santis, P. Mortier, M. De Beule, P. Segers, P. Verdonck, and B. Verhegghe, “Patient-specific Computational Fluid Dynamics: Structured mesh generation from coronary angiography,” Med. Biol. Eng. Comput., vol. 48, no. 4, pp. 371–380, 2010.

[21] A. Creane, E. Maher, S. Sultan, N. Hynes, D. J. Kelly, and C. Lally, “Finite element modelling of diseased carotid bifurcations generated from in vivo computerised tomographic angiography,” Comput. Biol. Med., vol. 40, no. 4, pp. 419–429, 2010.

[22] J. Ohayon, P. Teppaz, G. Finet, and G. Rioufol, “In-vivo prediction of human coronary plaque rupture location using intravascular ultrasound and the finite element method,” Coron. Artery Dis., vol. 12, no. 8, pp. 655–663, 2001.

[23] R. A. Baldewsing, C. L. De Korte, J. A. Schaar, F. Mastik, and A. F. W. Van der Steen, “Finite element modeling and intravascular ultrasound elastography of vulnerable plaques: Parameter variation,” Ultrasonics, vol. 42, no. 1–9, pp. 723–729, 2004.

[24] J. Tarjuelo-Gutierrez et al., “High-quality conforming hexahedral meshes of patient-specific abdominal aortic aneurysms including their intraluminal thrombi,” Med. Biol. Eng. Comput., vol. 52, no. 2, pp. 159–168, 2014.

[25] X. Huang, Z. Teng, G. Canton, M. Ferguson, C. Yuan, and D. Tang, “Intraplaque hemorrhage is associated with higher structural stresses in human atherosclerotic plaques: An in vivo MRI-based 3d fluid-structure interaction study,” Biomed. Eng. Online, vol. 9, pp. 1–12, 2010.

[26] Y. Zhang, Y. Bazilevs, S. Goswami, C. L. Bajaj, and T. J. R. Hughes, “Patient-specific vascular NURBS modeling for isogeometric analysis of blood flow,” Proc. 15th Int. Meshing Roundtable, IMR 2006, vol. 196, pp. 73–92, 2006.

[27] B. J. B. M. Wolters, M. C. M. Rutten, G. W. H. Schurink, U. Kose, J. De Hart, and F. N. Van De Vosse, “A patient-specific computational model of fluid-structure interaction in abdominal aortic aneurysms,” Med. Eng. Phys., vol. 27, no. 10, pp. 871–883, 2005.

[28] J. Bols et al., “Unstructured hexahedral mesh generation of complex vascular trees using a multi-block grid-based approach,” Comput. Methods Biomech. Biomed. Engin., vol. 19, no. 6, pp. 663–672, 2016.

[29] J. Bols et al., “Unstructured hexahedral mesh generation of complex vascular trees using a multi-block grid-based approach,” Comput. Methods Biomech. Biomed. Engin., vol. 19, no. 6, pp. 663–672, 2016.

[30] M. L. Raghavan, B. Ma, and M. F. Fillinger, “Non-invasive determination of zero-pressure geometry of arterial aneurysms,” Ann. Biomed. Eng., vol. 34, no. 9, pp. 1414–1419, 2006.

[31] F. Riveros, S. Chandra, E. A. Finol, T. C. Gasser, and J. F. Rodriguez, “A pull-back algorithm to determine the unloaded vascular geometry in anisotropic hyperelastic AAA passive mechanics,” Ann. Biomed. Eng., vol. 41, no. 4, pp. 694–708, 2013.

[32] M. L. Raghavan, B. Ma, and M. F. Fillinger, “Non-invasive determination of zero-pressure geometry of arterial aneurysms,” Ann. Biomed. Eng., vol. 34, no. 9, pp. 1414–1419, 2006.

[33] J. Bols, J. Degroote, B. Trachet, B. Verhegghe, P. Segers, and J. Vierendeels, “A computational method to assess the in vivo stresses and unloaded configuration of patient-specific blood vessels,” J. Comput. Appl. Math., vol. 246, pp. 10–17, 2013.

[34] S. de Putter, B. J. B. M. Wolters, M. C. M. Rutten, M. Breeuwer, F. A. Gerritsen, and F. N. van de Vosse, “Patient-specific initial wall stress in abdominal aortic aneurysms with a backward incremental method,” J. Biomech., vol. 40, no. 5, pp. 1081–1090, 2007.

[35] C. J. Chuong and Y. C. Fung, “Three-dimensional stress distribution in arteries.,” J. Biomech. Eng., vol. 105, no. 3, pp. 268–74, 1983.

[36] C. J. Chuong and Y. C. Fung, “On Residual Stresses in Arteries,” J. Biomech. Eng., vol. 108, no. May, pp. 189–192, 1986.

[37] Y. C. Fung, Biomechanics: Mechanical Properties of Living Tissues, Second. Media, Springer Science & Business, 1981.

[38] Y. C. Fung, “What Are the Residual Stresses Doing in Our Blood Vessels?,” Ann. Biomed. Eng., vol. 19, pp. 237–249, 1991.

[39] A. Delfino, N. Stergiopulos, J. E. Moore, and J. J. Meister, “Residual strain effects on the stress field in a thick wall finite element model of the human carotid bifurcation,” J. Biomech., vol. 30, no. 8, pp. 777–786, 1997.

[40] M. L. Raghavan, S. Trivedi, A. Nagaraj, D. D. McPherson, and K. B. Chandran, “Three-dimensional finite element analysis of residual stress in arteries,” Ann. Biomed. Eng., vol. 32, no. 2, pp. 257–263, 2004.

[41] R. N. Vaishnav and J. Vossoughi, “Residual stress and strain in aortic segments,” J. Biomech., vol. 20, no. 3, 1987.

[42] D. M. Pierce et al., “A method for incorporating three-dimensional residual stretches/stresses into patient-specific finite element simulations of arteries,” J. Mech. Behav. Biomed. Mater., vol. 47, pp. 147–164, 2015.

[43] G. A. Holzapfel and R. W. Ogden, “Modelling the layer-specific three-dimensional residual stresses in arteries, with an application to the human aorta,” J. R. Soc. Interface, vol. 7, no. 46, pp. 787–799, 2010.

[44] V. Alastrué, E. Peña, M.Á. Martínez, and M. Doblaré, “Numerical framework for patient-specific computational modelling of vascular tissue,” Int. j. numer. method. biomed. eng., vol. 26, no. 1, pp. 807–827, 2010.

[45] V. Alastrué, E. Peña, M.Á. Martínez, and M. Doblaré, “Assessing the use of the ‘opening angle method’ to enforce residual stresses in patient-specific arteries,” Ann. Biomed. Eng., vol. 35, no. 10, pp. 1821–1837, 2007.

[46] S. Saini, A., Berry, C., & Greenwald, “Effect of Age and Sex on Residual Stress in the Aorta,” J. Vasc. Res., vol. 32, no. 6, p. 43, 1995.

[47] J. Urevc, M. Halilovic, M. Brumen, and B. Štok, “An approach to consider the arterial residual stresses in modelling of a patient-specific artery,” Adv. Mech. Eng., vol. 8, no. 11, pp. 1–19, 2016.

[48] G. A. H. Hannah Weisbecker, David M. Pierce, “A generalized prestressing algorithm for finite element simulations of preloaded geometries with application to the aorta,” Int. j. numer. method. biomed. eng., vol. 26, no. 1, pp. 807–827, 2010.

[49] J. Schröder and S. Brinkhues, “A novel scheme for the approximation of residual stresses in arterial walls,” Arch. Appl. Mech., vol. 84, no. 6, pp. 881–898, 2014.

[50] S. Avril, P. Badel, and A. Duprey, “Anisotropic and hyperelastic identification of in vitro human arteries from full-field optical measurements,” J. Biomech., vol. 43, no. 15, pp. 2978–2985, 2010.

[51] K. Genovese, Y. U. Lee, A. Y. Lee, and J. D. Humphrey, “An improved panoramic digital image correlation method for vascular strain analysis and material characterization,” J. Mech. Behav. Biomed. Mater., vol. 27, pp. 132–142, 2013.

[52] M. Kroon and G. A. Holzapfel, “Estimation of the Distributions of Anisotropic, Elastic Properties and Wall Stresses of Saccular Cerebral Aneurysms by Inverse Analysis Estimation elastic of the distributions and wall properties cerebral aneurysms by of anisotropic, of saccular analysi,” Proc. R. Soc., vol. 464, no. 2092, pp. 807–825, 2008.

[53] A. C. Akyildiz et al., “A Framework for Local Mechanical Characterization of Atherosclerotic Plaques: Combination of Ultrasound Displacement Imaging and Inverse Finite Element Analysis,” Ann. Biomed. Eng., vol. 44, no. 4, pp. 968–979, 2016.

[54] A. Pandit, X. Lu, C. Wang, and G. S. Kassab, “Biaxial elastic material properties of porcine coronary media and adventitia,” Am. J. Physiol. Circ. Physiol., vol. 288, no. 6, pp. H2581– H2587, 2005.

[55] M. Ghasemi, D. R. Nolan, and C. Lally, “An investigation into the role of different constituents in damage accumulation in arterial tissue and constitutive model development,” Biomech. Model. Mechanobiol., no. i, 2018.

[56] G. Sommer, P. Regitnig, L. Költringer, and G. A. Holzapfel, “Biaxial mechanical properties of intact and layer-dissected human carotid arteries at physiological and supraphysiological loadings,” Am. J. Physiol. Circ. Physiol., vol. 298, no. 3, pp. H898–H912, 2009.

[57] R. D. Johnston, R. T. Gaul, and C. Lally, “An investigation into the critical role of fibre orientation in the ultimate tensile strength and stiffness of human carotid plaque caps,” Acta Biomater., no. xxxx, 2021.

[58] Z. Teng, D. Tang, J. Zheng, P. K. Woodard, and A. H. Hoffman, “An experimental study on the ultimate strength of the adventitia and media of human atherosclerotic carotid arteries in circumferential and axial directions,” J. Biomech., vol. 42, no. 15, pp. 2535–2539, 2009.

[59] Z. Teng et al., “The influence of constitutive law choice used to characterise atherosclerotic tissue material properties on computing stress values in human carotid plaques,” J. Biomech., vol. 48, no. 14, pp. 3912–3921, 2015.

[60] M. Liu, L. Liang, and W. Sun, “A new inverse method for estimation of in vivo mechanical properties of the aortic wall,” J. Mech. Behav. Biomed. Mater., vol. 72, no. January, pp. 148– 158, 2017.

[61] A. Wittek et al., “A fi nite element updating approach for identi fi cation of the anisotropic hyperelastic properties of normal and diseased aortic walls from 4D ultrasound strain imaging,” J. Mech. Behav. Biomed. Mater., vol. 58, pp. 122–138, 2016.

[62] A. Wittek et al., “In vivo determination of elastic properties of the human aorta based on 4D ultrasound data,” J. Mech. Behav. Biomed. Mater., vol. 27, pp. 167–183, 2013.

[63] K. Miller and J. Lu, “On the prospect of patient-specific biomechanics without patient-specific properties of tissues.,” J. Mech. Behav. Biomed. Mater., vol. 27, pp. 154–66, 2013.

[64] G. A. Holzapfel, T. C. Gasser, and R. A. Y. W. Ogden, “A New Constitutive Framework for Arterial Wall Mechanics and a Comparative Study of Material Models,” pp. 1–48, 2001.

[65] D. Balzani, S. Brinkhues, and G. A. Holzapfel, “Constitutive framework for the modeling of damage in collagenous soft tissues with application to arterial walls,” Comput. Methods Appl. Mech. Eng., vol. 213–216, pp. 139–151, 2012.

[66] H. A. Nieuwstadt et al., “A computer-simulation study on the effects of MRI voxel dimensions on carotid plaque lipid-core and fibrous cap segmentation and stress modeling,” PLoS One, vol. 10, no. 4, pp. 1–15, 2015.

