## Supplementary document for "Inverse Material Parameter Estimation of Patient Specific Finite Element Models at the Carotid Bifurcation: The Impact of Excluding the Zero Pressure Configuration and Residual Stress"

**Table 1: Material parameters used for each case for simulation of the stress for geometry 2. Case 2, 3 and 4 have been optimized under certain conditions previously stated in the text**

| Parameters | $\boldsymbol{C}_{\boldsymbol{10}}$ *(kPa)* | $\boldsymbol{k}_{\boldsymbol{1}}$ *(kPa)* | $\boldsymbol{k}_{\boldsymbol{2}}$ | $\boldsymbol{\kappa}$ | $\boldsymbol{\theta}$ *(deg)* |
| --- | --- | --- | --- | --- | --- |
| Case 1 | 6.56 | 1482 | 561 | 0.16 | 37 |
| Case 2 | 5.82 | 1512 | 522 | 0.16 | 37 |
| Case 3 | 8.35 | 1428.61 | 941.13 | 0.12 | 38 |
| Case 4 | 9.42 | 1489.1 | 385.61 | 0.20 | 37 |

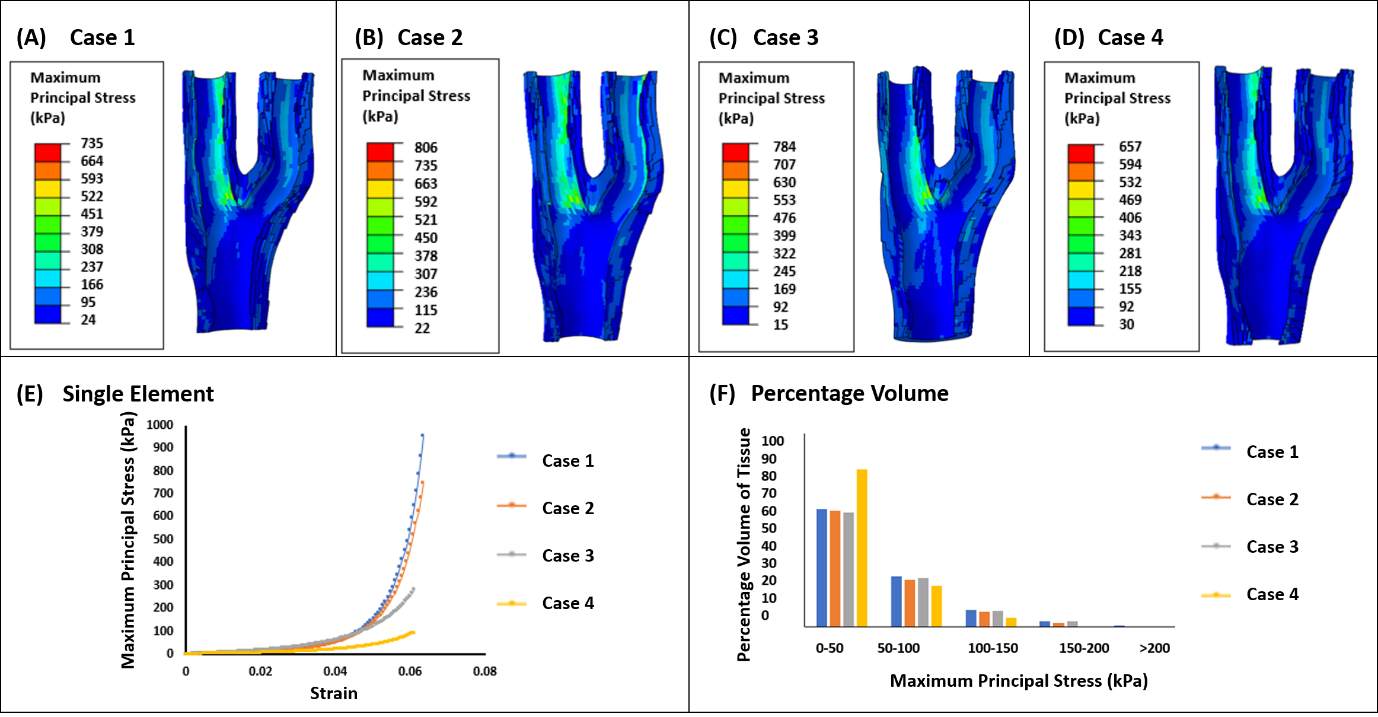

**Figure A1: Maximum Principal Stress (kPa) observed in geometry 2 with (i) material parameters taken from literature (Case 1) (ii) optimized parameters without the inclusion of zero pressure configuration and residual stress (Case 2) (iii) optimized parameters with the inclusion of the zero pressure configuration (Case 3) (iv) optimized parameters with the inclusion of the zero pressure configuration and residual stress (Case 4) (v) Single element test undergoing uniaxial tension looking at stress strain response of each case (vi) Percentage of volume graph for the 4 cases showing the stress distribution throughout the vessel wall**

**Table 2: Material parameters used for each case for simulation of the stress for geometry 3. Case 2, 3 and 4 have been optimized under certain conditions previously stated in the text**

| Parameters | $\boldsymbol{C}_{\boldsymbol{10}}$ *(kPa)* | $\boldsymbol{k}_{\boldsymbol{1}}$ *(kPa)* | $\boldsymbol{k}_{\boldsymbol{2}}$ | $\boldsymbol{\kappa}$ | $\boldsymbol{\theta}$ *(deg)* |
| --- | --- | --- | --- | --- | --- |
| Case 1 | 6.56 | 1482 | 561 | 0.16 | 37 |
| Case 2 | 3.26 | 1472 | 571 | 0.16 | 37 |
| Case 3 | 6.06 | 1479.29 | 862.02 | 0.17 | 39 |
| Case 4 | 8.68 | 1546.92 | 423.17 | 0.22 | 39 |

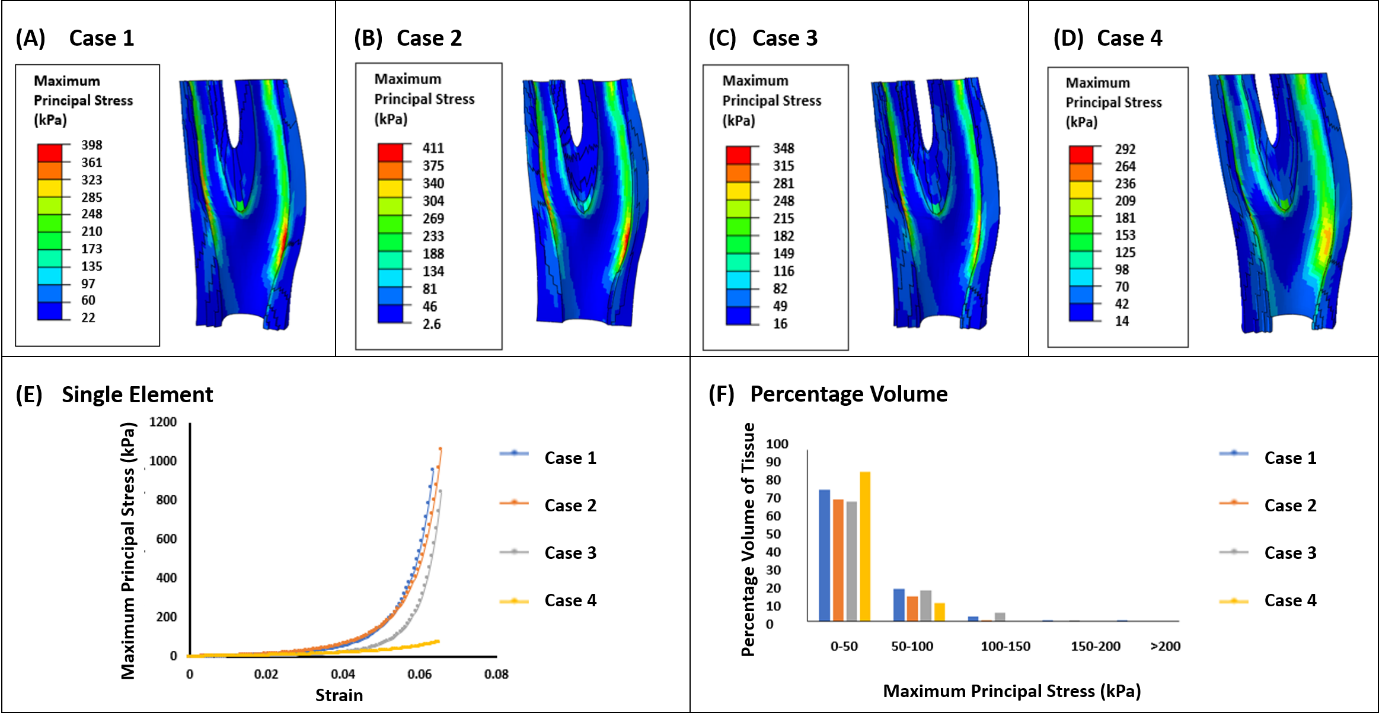

**Figure A2: Maximum Principal Stress (kPa) observed in geometry 3 with (i) material parameters taken from literature (Case 1) (ii) optimized parameters without the inclusion of zero pressure configuration and residual stress (Case 2) (iii) optimized parameters with the inclusion of the zero pressure configuration (Case 3) (iv) optimized parameters with the inclusion of the zero pressure configuration and residual stress (Case 4) (v) Single element test undergoing uniaxial tension looking at stress strain response of each case (vi) Percentage of volume graph for the 4 cases showing the stress distribution throughout the vessel wall**

**Table 3: Material parameters used for each case for simulation of the stress for geometry 4. Case 2, 3 and 4 have been optimized under certain conditions previously stated in the text**

| Parameters | $\boldsymbol{C}_{\boldsymbol{10}}$ *(kPa)* | $\boldsymbol{k}_{\boldsymbol{1}}$ *(kPa)* | $\boldsymbol{k}_{\boldsymbol{2}}$ | $\boldsymbol{\kappa}$ | $\boldsymbol{\theta}$ *(deg)* |
| --- | --- | --- | --- | --- | --- |
| Case 1 | 6.56 | 1482 | 561 | 0.16 | 37 |
| Case 2 | 8.82 | 1568 | 586 | 0.16 | 37 |
| Case 3 | 7.98 | 1653.13 | 616.09 | 0.15 | 38 |
| Case 4 | 9.64 | 1369.46 | 224.64 | 0.24 | 38 |

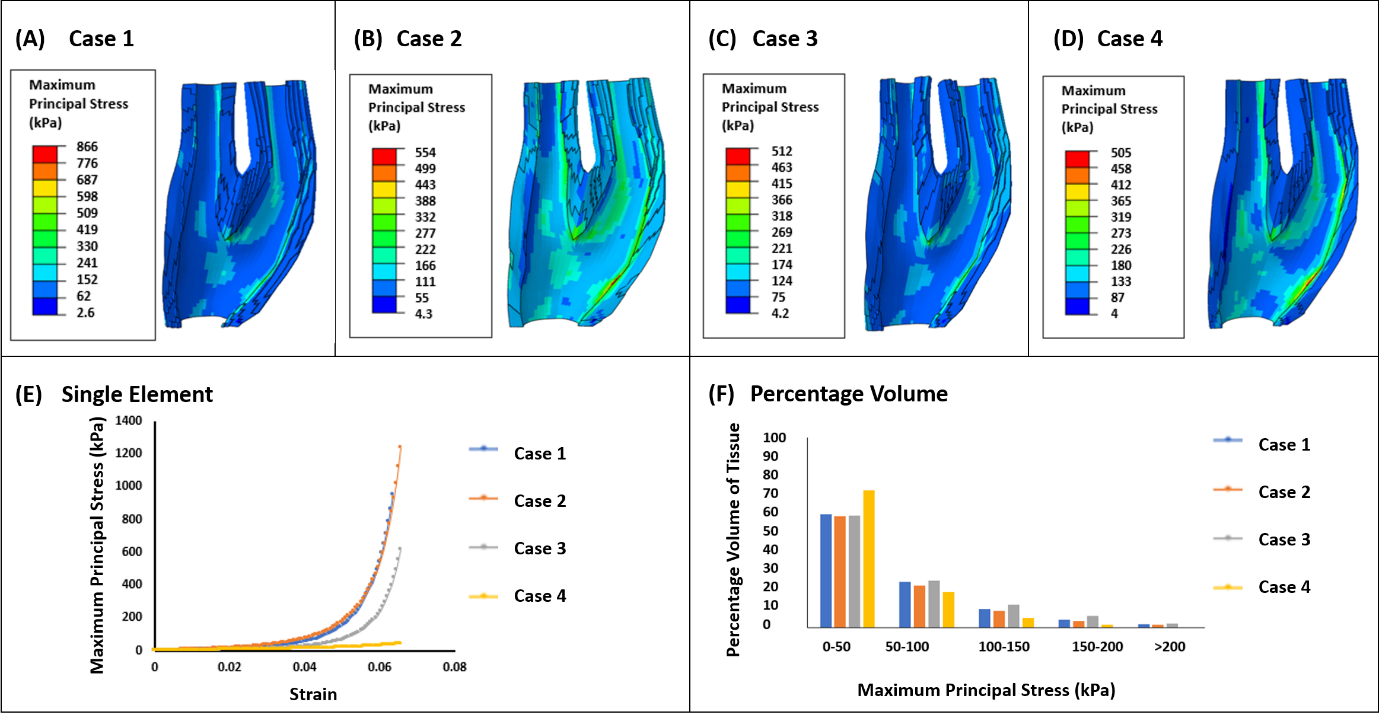

**Figure A3: Maximum Principal Stress (kPa) observed in geometry 4 with (i) material parameters taken from literature (Case 1) (ii) optimized parameters without the inclusion of zero pressure configuration and residual stress (Case 2) (iii) optimized parameters with the inclusion of the zero pressure configuration (Case 3) (iv) optimized parameters with the inclusion of the zero pressure configuration and residual stress (Case 4) (v) Single element test undergoing uniaxial tension looking at stress strain response of each case (vi) Percentage of volume graph for the 4 cases showing the stress distribution throughout the vessel wall**

**Table 4: Material parameters used for each case for simulation of the stress for geometry 5. Case 2, 3 and 4 have been optimized under certain conditions previously stated in the text**

| Parameters | $\boldsymbol{C}_{\boldsymbol{10}}$ *(kPa)* | $\boldsymbol{k}_{\boldsymbol{1}}$ *(kPa)* | $\boldsymbol{k}_{\boldsymbol{2}}$ | $\boldsymbol{\kappa}$ | $\boldsymbol{\theta}$ *(deg)* |
| --- | --- | --- | --- | --- | --- |
| Case 1 | 6.56 | 1482 | 561 | 0.16 | 37 |
| Case 2 | 12.46 | 1549 | 561 | 0.16 | 37 |
| Case 3 | 7.75 | 1722.37 | 684.81 | 0.12 | 37 |
| Case 4 | 8.76 | 1682.12 | 464.21 | 0.18 | 38 |

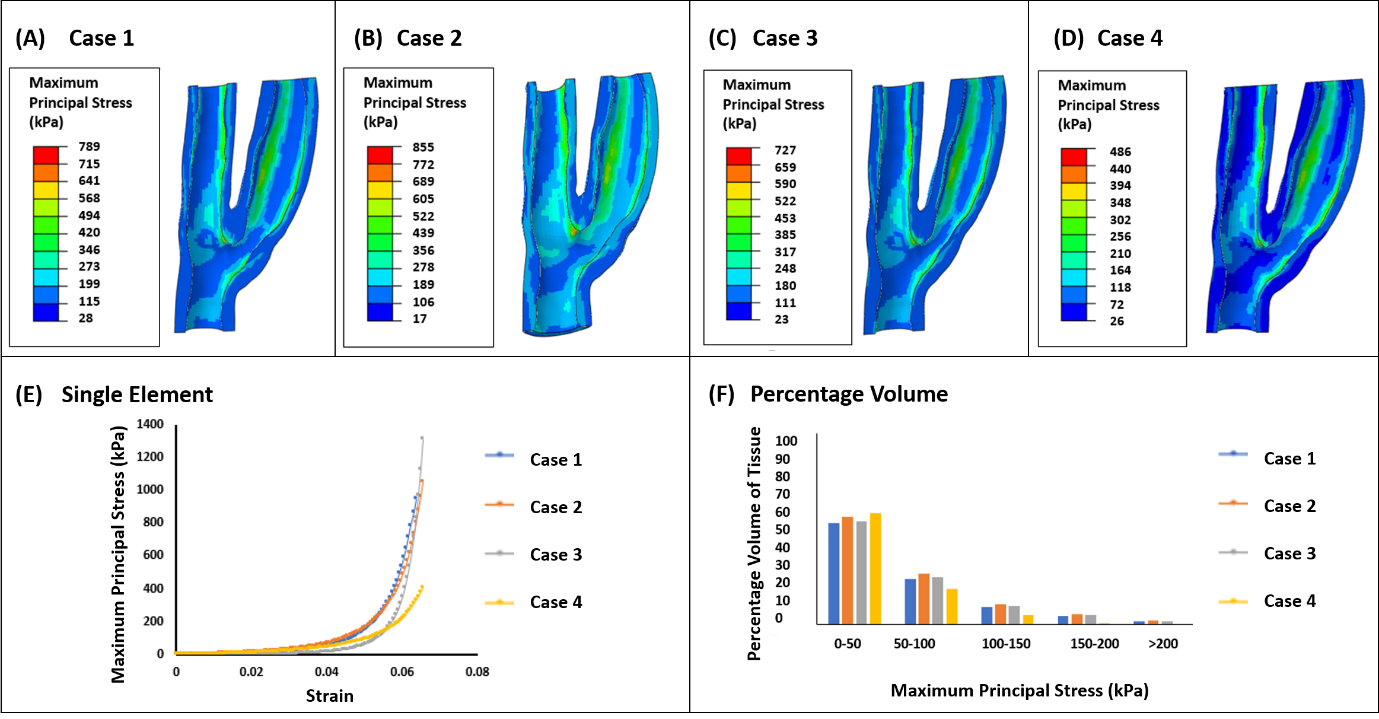

**Figure A4: Maximum Principal Stress (kPa) observed in geometry 5 with (i) material parameters taken from literature (Case 1) (ii) optimized parameters without the inclusion of zero pressure configuration and residual stress (Case 2) (iii) optimized parameters with the inclusion of the zero pressure configuration (Case 3) (iv) optimized parameters with the inclusion of the zero pressure configuration and residual stress (Case 4) (v) Single element test undergoing uniaxial tension looking at stress strain response of each case (vi) Percentage of volume graph for the 4 cases showing the stress distribution throughout the vessel wall**

**Table 5: Material parameters used for each case for simulation of the stress for geometry 6. Case 2, 3 and 4 have been optimized under certain conditions previously stated in the text**

| Parameters | $\boldsymbol{C}_{\boldsymbol{10}}$ *(kPa)* | $\boldsymbol{k}_{\boldsymbol{1}}$ *(kPa)* | $\boldsymbol{k}_{\boldsymbol{2}}$ | $\boldsymbol{\kappa}$ | $\boldsymbol{\theta}$ *(deg)* |
| --- | --- | --- | --- | --- | --- |
| Case 1 | 6.56 | 1482 | 561 | 0.16 | 37 |
| Case 2 | 6.87 | 1362 | 486 | 0.16 | 37 |
| Case 3 | 6.43 | 1280.77 | 744.01 | 0.08 | 38 |
| Case 4 | 10.24 | 1864.22 | 486.46 | 0.18 | 37 |

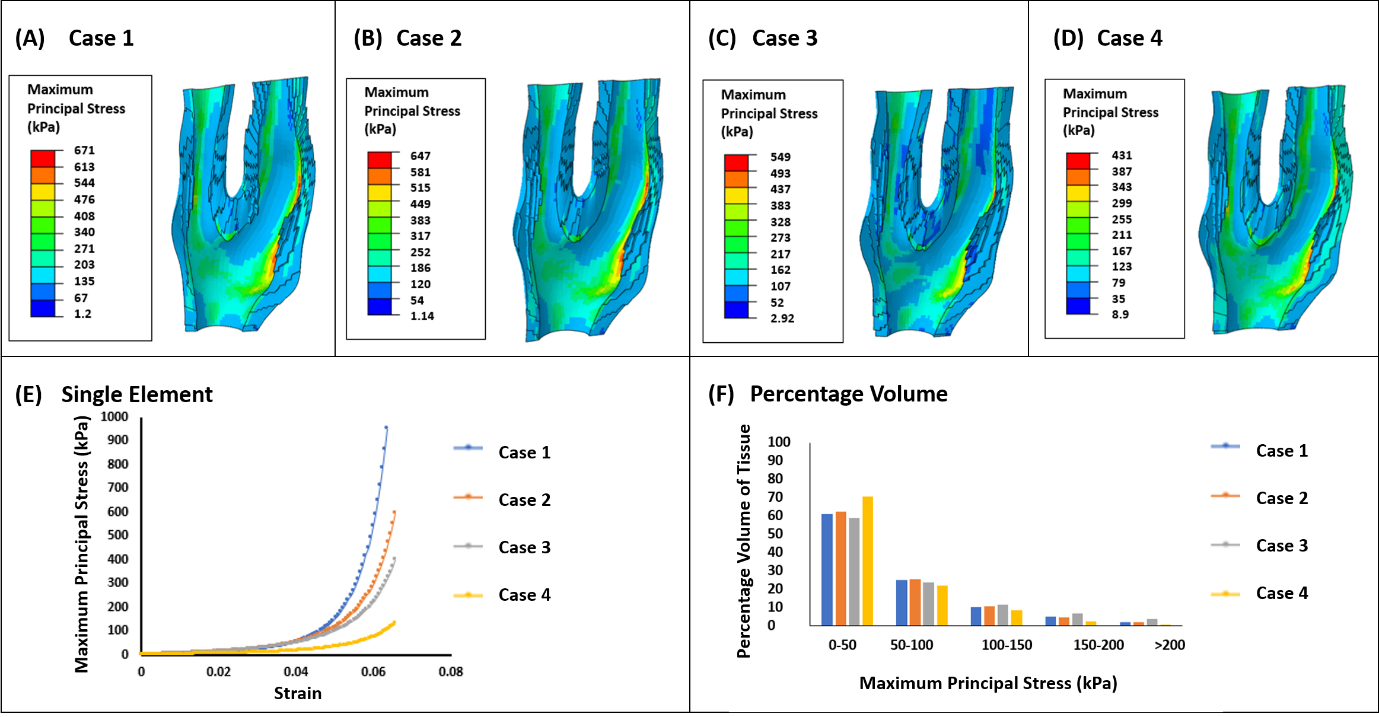

**Figure A5: Maximum Principal Stress (kPa) observed in geometry 6 with (i) material parameters taken from literature (Case 1) (ii) optimized parameters without the inclusion of zero pressure configuration and residual stress (Case 2) (iii) optimized parameters with the inclusion of the zero pressure configuration (Case 3) (iv) optimized parameters with the inclusion of the zero pressure configuration and residual stress (Case 4) (v) Single element test undergoing uniaxial tension looking at stress strain response of each case (vi) Percentage of volume graph for the 4 cases showing the stress distribution throughout the vessel wall**
